# Active Escape or Active Change? Decomposing Pavlovian Bias into Non-emotional Components

**DOI:** 10.64898/2026.09.17.751801

**Authors:** Yi-An A. Chen, Yu-Fan Wu, Yu-Wen Jan, Shi-Jen Tsai

## Abstract

Pavlovian bias, a motor bias driven by emotionally-valenced context, has long been considered a proxy for emotion-action processing, which may become aberrant in affective disorders and suicidality. The orthogonal valenced Go/NoGo paradigm was developed to isolate this bias and has been widely adopted in translational psychiatry research under this interpretation. However, several unresolved theoretical inconsistencies highlight the need to dissociate potential confounding factors within this paradigm before assigning mechanistic and clinical interpretations to its findings. Using a novel paradigm that fully orthogonalized valence, action, and aimed transition in a normative sample of 25 individuals, we found that the “active escape bias” previously observed in aversive contexts—and interpreted as reflecting suicidality in the clinical population—was reproduced in appetitive contexts using trials with the same transition structure. Furthermore, computational modeling indicated that the bias patterns isolated by the orthogonal paradigm are primarily explained by the combined effects of aimed transition and cue salience, without requiring emotional-valence modulation. These findings call for a fundamental reconsideration of how orthogonal Go/NoGo paradigms are interpreted in translational psychiatry, particularly how the behavior they isolate is mechanistically linked to affective symptoms.

## Introduction

Human decision-making is inherently imperfect. Prime examples are various forms of biases, which can override objective evaluations of circumstances and lead to suboptimal choices [1, 2]. While common, severe and persistent biases—such as selective processing of negative events—can signal aberrant neural processes linked to psychiatric disorders [3, 4]. Therefore, many behavioral paradigms have been developed to examine how fundamental forms of biases relate to psychiatric symptoms [5–7].

Pavlovian bias, defined as a motor bias introduced by emotional cues, has been extensively investigated in cognitive neuropsychiatry due to its potential link to affective disorders [5, 8–11]. One paradigm for examinining such bias, originally proposed by Guitart-Masip et al. [12], leverages orthogonal action (Go v.s. NoGo) and valence (reward v.s. punishment) pairings to expose biased associations between emotions and motor responses that can override objective reinforcement learning. Their study demonstrated an inherent bias to emit a response (i.e. Go) to obtain reward alongside a bias to withhold a response (i.e. NoGo) to avoid punishment, both interpreted as learned emotional context (appetitive for reward and aversive for punishment) biasing action selection.

However, this version of Go/NoGo paradigm does not account for “active avoidance,” the bias toward executing a response under threat. Active avoidance is well-documented in conventional Pavlovian-to-Instrumental Transfer (PIT) paradigms and aligns with the classic fight-or-flight theories of stress [5, 8, 13, 14]. To address this gap, Millner et al. adapted the paradigm by substituting the reward-punishment axis with a threat-immediacy axis, converting reward-obtaining trials to threat-escaping trials [15]. Their study revealed a Go-bias on threat-escaping trials comparable to that observed on reward-obtaining trials. Drawing on the two-factor theory [13, 16, 17], the authors suggested this “active escape bias” is driven by (1) the learned association between the escape cue and “relief/safety,” which functions similarly to a reward, and (2) the intrinsic aversiveness of the escape cue. Subsequent studies have utilized this paradigm to investigate suicidal behavior, framing the active escape bias as a tendency to act upon negative emotions that mirrors suicidal intent [10, 18, 19].

Two critical issues emerge from the interpretation. First, the two-factor theory implies a gradual valence reversal of the escape cue (i.e. shifting from signaling threat to signaling relief/safety), a process that is difficult to test empirically given its latent and subjective nature. Second, the account cannot explain why the unconditioned aversiveness of escape cue induces a Go-bias, whereas the conditioned aversiveness of avoid cue induces a NoGo-bias. While these concerns might appear purely theoretical, they directly shape how the target symptoms (e.g. suicidal behavior) are interpreted and how clinical populations are conceptualized. They should therefore be approached with caution.

To develop a more parsimonious explanatory framework, we reviewed the trial structures of Guitart-Masip et al. and Millner et al. (Fig. 1, Obtain, Avoid and Escape) to identify factors that might have been overlooked [12, 15]. As illustrated in Fig. 1, a key distinction naturally emerges and sets the Avoid trial apart from the other two trial types (Obtain and Escape) when viewed through the lens of “aimed transitions.” Specifically, while all three trial types have a “better” outcome and a “worse” outcome, only the Avoid trial achieves the better outcome by “staying unchanged” (i.e. transitioning from a neutral cue to a neutral outcome). Conversely, the Obtain and Escape trials require a change: the former transitions from a neutral cue to a good outcome, whereas the latter transitions from a bad cue to a neutral outcome. The aimed transitions—“stay” versus “change”—could account for the observed pattern of motor bias.

**Figure 1:**
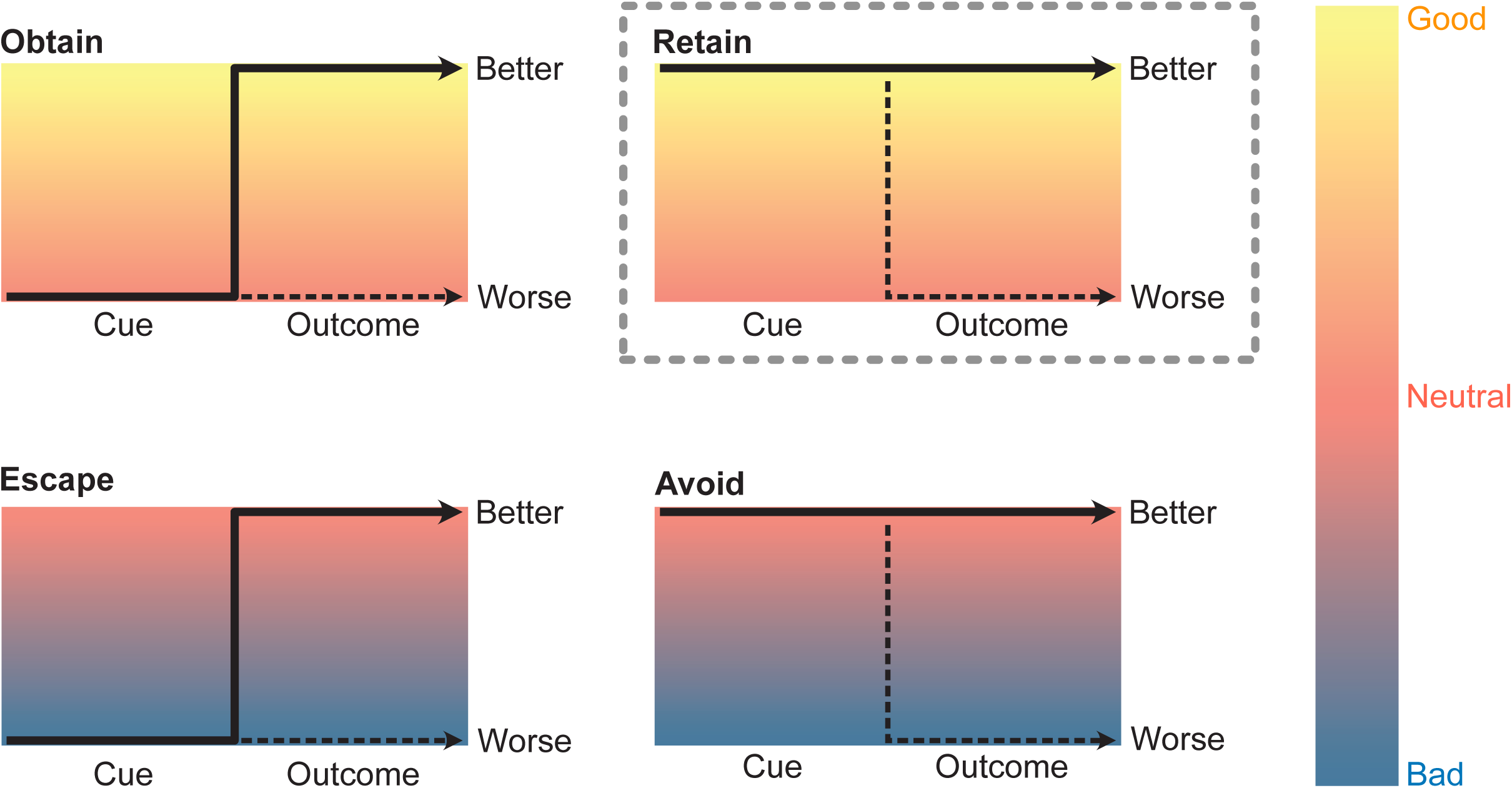
Illustration of the three trial structures from prior work alongside the novel trial structure (Retain) introduced in the present study. The absolute emotional valence associated with each trial—bad, neutral or good—is indicated by the background color underlying each trial. Solid arrows denote the intended transition leading to the better outcome, while dashed arrows denote the unintended transition leading to the worse outcome. As illustrated, “better” and “worse” are defined relative to each trial context (e.g. neutral is “worse” in the Obtain trial but “better” in the Escape trial). Reward and punishment trials in Guitart-Masip et al. correspond to the Obtain and Avoid structures [12], whereas escape and avoid trials in Millner et al. correspond to the Escape and Avoid structures [15]. The novel Retain structure introduced in the current study completes a fully orthogonalized factorial design crossing context valence with transitional structure.

An intuitive real-world analog of this transition-related bias can be seen in baseball. When facing an incoming pitch, batters naturally swing if they want to alter its trajectory and withhold the swing if they want it to pass undisturbed. Although alternative outcomes are possible (e.g. swinging and missing, or the ball striking a stationary bat), we automatically map “swinging” onto “changing” and “withholding” onto “staying unchanged.” Similarly, the Go-bias on Escape and Obtain trials may stem from an intrinsic link between “action” and “changing”, whereas the NoGo-bias on Avoid trial may reflect a link between “inaction” and “staying unchanged.” Unlike the two-factor theory, the hypothesis of transition-related bias is grounded in intuitive expectations about physical laws and does not require a valence reversal.

Another distinction is the presence of non-neutral cue, which sets the Escape trial apart from the other two trial types (Obtain and Avoid; Fig. 1). While Millner et al. suggested that active escape bias could arise from the “aversiveness” of the Escape cue [15], cue non-neutrality itself—whether aversive or appetitive—may be sufficient to drive motor bias due to its salience relative to baseline [20–22]. Owing to design limitations, the Obtain trial cannot incorporate a non-neutral cue to enable a direct comparison with its aversive counterpart (i.e. the Escape trial). Nevertheless, a non-neutral appetitive cue can be accommodated by a trial type featuring a “stay” transition (Retain; upper right of Fig. 1). The inclusion of the novel Retain trial completes a fully orthogonalized 2 ✕ 2 factorial design crossing valence (appetitive vs. aversive) and transition (change vs. stay). By further incorporating the action choices (Go vs. NoGo), this 2 ✕ 2 ✕ 2 framework enables the precise dissociation of transition-related and valence-related biases.

Based on this factorial design, we developed a novel behavioral paradigm in which participants maximize gains and minimize losses by learning action-outcome contingencies across four contexts featuring distinct valence/action-congruency combinations. We aim to address two core questions: (1) Can the bias pattern in the aversive learning context (i.e. Escape and Avoid) reported by Millner et al. be replicated in an appetitive learning context sharing identical transition structures (i.e. Obtain and Retain)? (2) Does an appetitive cue (i.e. Retain cue) bias motor response in the same manner as an aversive cue (i.e. Escape cue)? We found that the bias pattern in the appetitive context closely mirrors that observed in the aversive context, and that the modest context-dependent difference is accounted for by cue salience, which invigorates actions similarly across both contexts. These findings indicate that the Pavlovian bias isolated by valenced orthogonal Go/NoGo paradigms may be largely attributed to two non-emotional components: aimed transition and cue salience. They therefore invite a reconsideration of how the Pavlovian bias—traditionally viewed as emotion-driven—should be interpreted in research on affective disorders and suicidality.

## Methods

### Participants

25 healthy participants aged 20–50 years (14 females; mean age = 34.96 ± 6.30) were recruited from local Taiwanese communities via public advertisements between May and July 2026. Exclusion criteria were: (1) a history of neurological disorders (e.g. epilepsy, stroke, or traumatic brain injury); (2) a history of neurodevelopmental, affective, psychotic, or personality disorders; and (3) a history of substance use disorders. After providing written informed consent, participants received a unique, one-time access link to complete the web-based task at their convenience via Cognition.run (https://www.cognition.run). Participants were compensated with 200 NTD upon task completion. The study protocol was approved by the Institutional Review Board of Taipei Veterans General Hospital (IRB #: 2026-03-003C).

We originally targeted a minimum sample size of 20 participants. To account for higher attrition and data quality variability in web-based experiments [23], we assumed a 25% data exclusion rate and recruited 28 participants. Of the 28 registered individuals, two were lost to follow-up after providing informed consent, and one withdrew midway through the task. For the 25 participants who completed the task, we adopted a data quality threshold of 55% accuracy. Because all 25 completers exceeded this threshold (mean accuracy = 73% ± 10%, range = [58%, 96%]), no data were excluded, yielding a final analytic sample of 25 participants.

### Task

Fig. 2a illustrates the eight trial conditions, each defined by pairing one of the four transition–valence combinations (Fig. 1) with a required action (Go vs. NoGo). In the Obtain trials, the cue displays a static image of an open bag, and the participant selects a response—either pressing or withholding a button press—to trigger gem inflow into the bag. In the Retain trials, the cue presents an animation of gems actively flowing into the bag, and the participant selects a response to extend this inflow. In the Escape trials, the cue presents an animation of gems leaking out through a break in the bag’s side, and the participant selects a response to stop the leak. In the Avoid trials, the cue shows a static image of a bag with side stitching, and the participant selects a response to prevent the stitches from rupturing and causing gem leakage. Across all trials, participants were required to select and execute a response within a 3-second cue display window. Executing a button press (i.e. a Go response) immediately triggered the transition from cue to outcome. If no button press was detected within 3 seconds, it was recorded as a NoGo response, and the cue automatically timed out and transitioned to the corresponding outcome.

**Figure 2:**
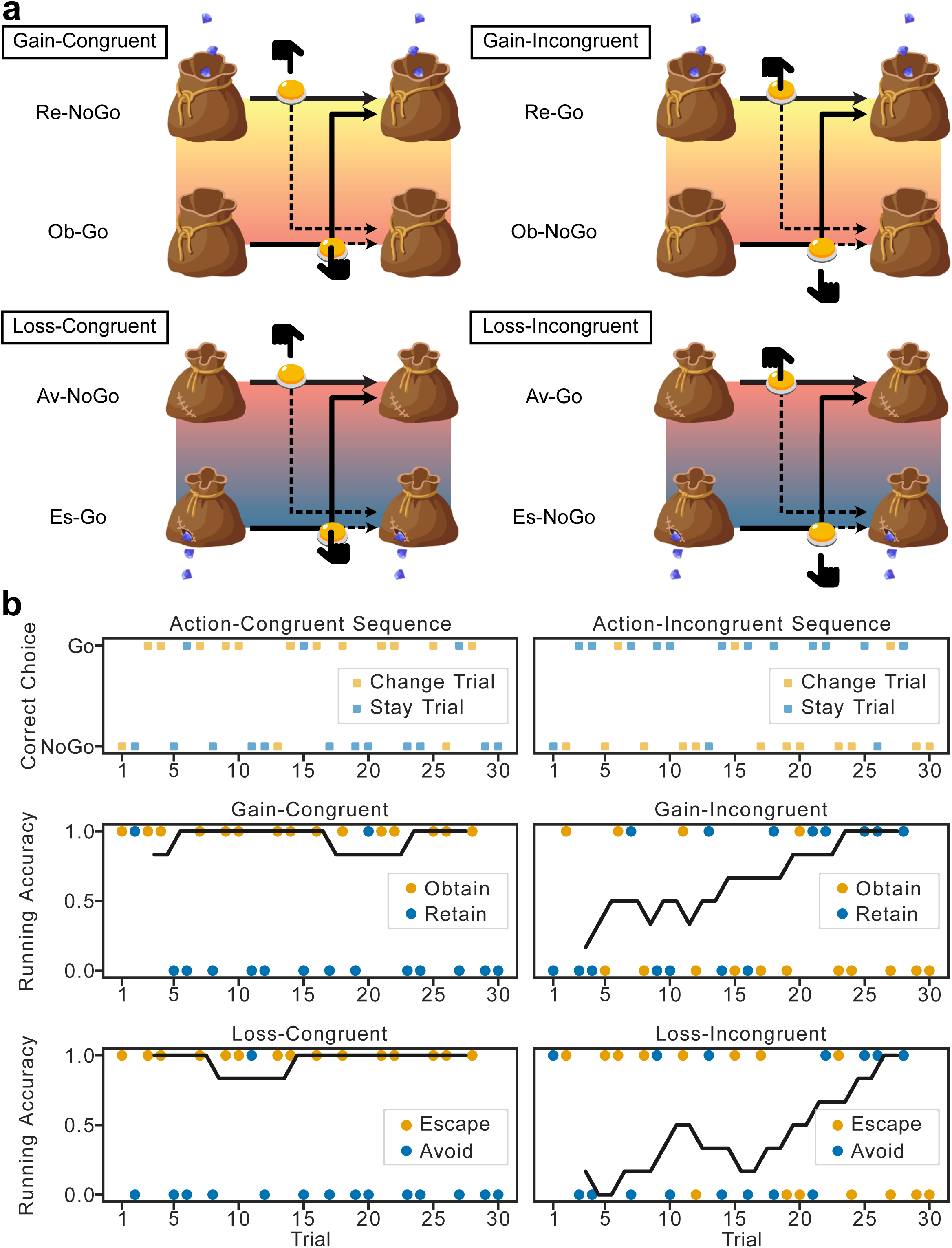
Task structure and representative participant data. (a) The four experimental blocks defined by context valence (Gain vs. Loss) and action congruency (Congruent vs. Incongruent). Each block comprises two dominant trial types, with cue phases displayed on the left and outcome phases on the right. Solid arrows denote intended transitions leading to better outcomes, whereas dashed arrows denote unintended transitions leading to worse outcomes. The button and hand icons indicate the required action—Go (finger pressing the button) or NoGo (finger not pressing the button)—to achieve the better outcome indicated by the solid arrow in each trial. Ob: Obtain; Re: Retain; Es: Escape; Av: Avoid. (b) Representative data from one participant. Choice sequences for congruent and incongruent blocks are shown in the top panels. Participant responses across the four blocks are presented in the bottom four panels. Note that “correct choice” in the top panels denotes trial-level correctness (i.e. defined by the immediate outcome received on each trial), whereas the overlaid 6-trial running accuracy curves in the bottom panels reflect block-level correctness (i.e. defined by block-wise action congruency).

As illustrated in Fig. 2a, the task comprised two gain blocks and two loss blocks. Each pair contained one action-congruent block and one action-incongruent block. Here, “congruency” reflects the hypothesized mapping between transition structure and action (i.e., change–Go, stay–NoGo). Specifically, in action-congruent blocks, pressing the button yielded an 80% probability of initiating a state change (e.g., triggering gem inflow or halting gem leakage), whereas withholding a response yielded an 80% probability of maintaining the current state (e.g., continuing gem inflow or preventing gem leakage). In action-incongruent blocks, these contingencies were reversed.

In the task, participants were instructed to maximize gem accumulation by increasing gem inflow and decreasing gem leakage. Prior to the main task, participants were prompted to complete a practice round comprising four 6-trial blocks to familiarize themselves with the task structure. Following a reminder that probabilistic contingencies could change at any time during the formal round, participants initiated the main task, which comprised four 30-trial blocks (15 “stay” trials, 15 “change” trials) and lasted approximately 17-20 minutes. Block order was counterbalanced across participants. For illustrative purposes, Fig. 2b displays representative data from one participant to demonstrate the task structure. The correct responses (i.e. the choice sequences) are shown in the top panels, and the responses made by the participant in each block are shown in the bottom panels, overlaid with 6-trial running accuracy curves.

### Computational Modelling

Three variants of reinforcement learning drift diffusion models (RL-DDM) were constructed and validated to investigate the cognitive mechanisms underlying the observed behavioral patterns. The mathematical formulas, validation, fitting, and comparison procedures are described below, while the rationale for each model specification is presented following the behavioral findings in the Results section.

#### RL-DDM: Basic Concept

The RL-DDM is a cognitive model that describes how beliefs about the values of available choices are updated through experience and how the continuously updated value estimates shape subsequent choices over time. A comprehensive review of RL-DDM is provided by Pedersen et al. [24], and a schematic overview of the DDM component is shown in Fig. 5a. We adopted this modelling framework to enable direct comparison with the work of Millner et al., which employed the same approach [15, 18].

In brief, RL-DDM comprises two components: a learning rule (the RL component), and a decision-making rule (the DDM component). The learning rule governs how choice values (Q-values) are updated based on experience, whereas the decision-making rule specifies how the updated Q-values are translated into the drift rate *V*, which, together with other DDM parameters, determines which choice is selected and the time required to make that choice (Supplementary Information, Part I).

For the RL component, we adopted the Rescorla-Wagner delta learning rule across all three model variants [25]:

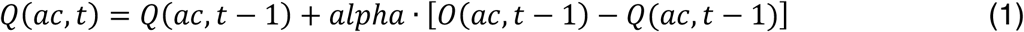

Here, *Q(ac, t)* and *Q(ac, t-1)*—both ranging from 0 to 1—represent the estimated value of choice *ac* at trials *t* and *t-1*, respectively. *O(ac, t-1)*—either 0 or 1—represents the outcome received at trial *t*-1 following choice *ac*. The parameter *alpha* is the learning rate, determining the extent to which new evidence is incorporated into the value estimate. In the present study, *ac* corresponds to “Go” or “NoGo”. For *O*, 0 represents the worse outcome and 1 represents the better outcome. The initial Q values of both action choices were set to 0.5, indicating that neither choice was assumed to be superior before any evidence was observed.

The updated Q values are translated into the drift rate in the DDM according to the following equation:

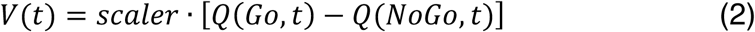

Here, *Q(Go, t)* and *Q(NoGo, t)* denote the estimated values of the Go and NoGo choices at trial *t*, respectively, whereas *scaler* is a positive scaling parameter that converts the Q-value difference into the drift rate at trial *t*, *V(t)*. As shown in Fig. 5a, the DDM requires two decision boundaries, with the upper boundary corresponding to Go and the lower boundary corresponding to NoGo in the present study. Consequently, a positive Q-value difference (i.e. *Q(Go, t)* > *Q(NoGo, t)*) produces a positive drift rate *V*, increasing the probability of reaching the Go boundary (i.e. pressing the button) while reducing the time required to reach it. Intuitively, as the difference between these two Q-values increases over trials through reinforcement learning, the agent becomes more confident about which choice is superior, resulting in faster and more deterministic decisions.

In this study, we built upon the findings of Millner et al. [15], adopting a parsimonious RL-DDM framework in which all model parameters were held constant across conditions except for the relative starting point *z*, which was modulated in a model variant-specific manner, as detailed below.

#### Model description

Three hierarchical Bayesian model variants were constructed using PyMC 6.2.0 to investigate different hypothesized mechanisms underlying the observed behavioral patterns [26]. The descriptions below focus on the modulation of the *z* parameter, which is the primary focus of the present study. For a detailed description of the priors and hyperpriors assigned to all RL-DDM parameters, please refer to part I of the Supplementary Information.

In Model 1 (M1), the relative starting point *z* is modulated by the aimed transition and cue salience according to the following equations:

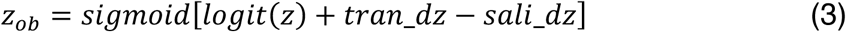

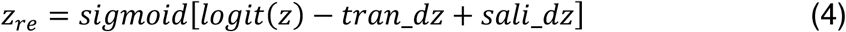

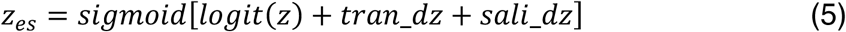

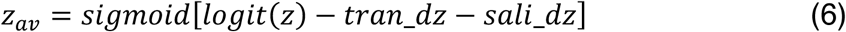

In equations (3)-(6), *z* denotes the baseline relative starting point, representing each participant’s baseline action tendency. A *z* value greater than 0.5 indicates an inherent tendency to press the button (i.e. a Go response) regardless of context, whereas a value less than 0.5 indicates an inherent tendency to withhold a button press (i.e. a NoGo response). This baseline starting point is further modulated by two parameters, *tran_dz* and *sali_dz*, which represent the shifts associated with the aimed transition and cue salience, respectively, across trial conditions. A positive *tran_dz* shifts the starting point toward the Go boundary on “change” trials and toward the NoGo boundary on “stay” trials. Likewise, a positive *sali_dz* shifts the starting point toward the Go boundary on trials with non-neutral cues and toward the NoGo boundary on trials with neutral cues. These modulations yield four condition-specific relative starting points: *z_ob_*, *z_re_*, *z_es_* and *z_av_*, corresponding to Obtain, Retain, Escape, and Avoid trials, respectively.

The sigmoid function and logit functions in equations (3)-(6) are mathematical inverses of each other:

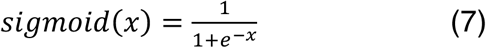

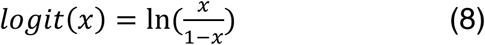

These transformations were required for *tran_dz* and *sali_dz* because both parameters were formed by normal priors centered at zero (Supplementary Information, Part I), reflecting the null hypothesis that they exert no effect on the relative starting point. Because these normal priors have unconstrained support, whereas *z* is constrained to the interval [0,1], the baseline *z* is first transformed to the unconstrained logit scale, where the additive modulation is applied. The resulting value is then transformed back to the bounded [0,1] scale using the sigmoid function.

In Model 2 (M2), the relative starting point *z* is modulated by the aimed transition and context valence according to the following equations:

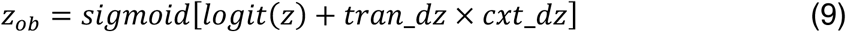

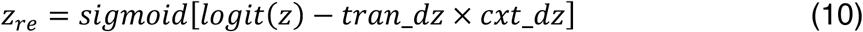

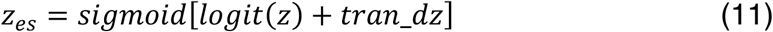

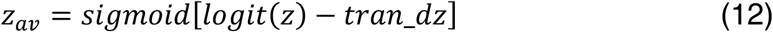

In M2, the action bias is hypothesized to be primarily driven by the aimed transition, with its magnitude further modulated by context valence. The direction of *tran_dz* remains identical to that in M1 across all trial conditions. A positive, log-normal-distributed scaling parameter with support on [0, ∞), *cxt_dz*, is applied multiplicatively to *tran_dz* on positively valenced trials (i.e. Obtain and Retain), thereby scaling the magnitude of the aimed-transition effect while preserving its direction in the appetitive context.

Model 3 (M3) is an exploratory extension from M1. In M3, the relative starting point *z* is modulated according to the following equations:

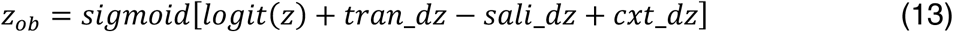

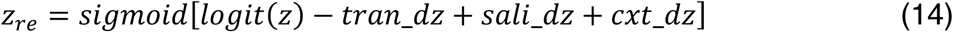

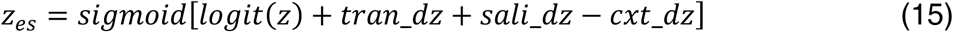

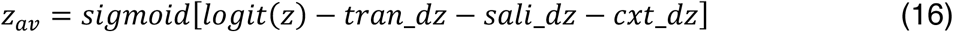

As shown in equations (13)-(16), and compared with the corresponding equations (3)-(6) for M1, M3 includes an additional additive modulation parameter, *cxt_dz*. This parameter shifts the relative starting point toward the Go boundary in positively valenced trials and towards the NoGo boundary in negatively valenced trials. Notably, *cxt_dz* is formulated differently in M2 and M3. In M3, *cxt_dz* functions as an additive modulation parameter, analogous to *tran_dz* and *sali_dz*, and therefore has a normal prior centered at zero with support over (-∞, ∞). In contrast, *cxt_dz* in M2 functions as a multiplicative modulation parameter with a log-normal prior centered at one and support over [0, ∞).

#### Model fitting procedure

To conduct Bayesian inference on *tran_dz*, *sali_dz* and *cxt_dz*, the three model variants were sampled using the No-U-Turn Sampler (NUTS) algorithm for Markov Chain Monte Carlo (MCMC) in PyMC 6.2.0 [27], with 1500 tuning steps, 4 MCMC chains, and 2000 posterior draws per chain. Convergence of the posterior samples was assessed using the Gelman-Rubin diagnostic (R^) and the effective sample size (ESS).

The log-likelihood function for MCMC sampling was built upon the Wiener First Passage Time (WFPT) distribution, a probability density function for choices and response times parameterized by drift rate *v*, boundary separation *a*, and relative starting point *z* [28, 29]. The standard form of WFPT distribution, which specifies the probability density of the response time *t* for reaching the lower boundary, is given below:

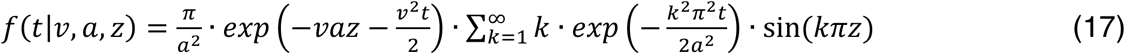

Equation (17) is used exclusively for trials with a Go response, as response times were recorded only for Go choices. Because Go choice was assigned to the upper boundary, *v* and *z* in equation (17) were replaced by –*v* and 1 – *z*, respectively, to evaluate the density at the upper boundary [28]. As equation (17) involves an infinite series that is computationally intractable, we utilized the approximation method proposed by Navarro and Fuss to compute the log-likelihood of Go responses [29].

For NoGo responses, which lack response time data, the log-likelihood was computed using the choice probability function, which specifies the marginal probability of reaching each decision boundary across all time. This function is obtained by integrating the WFPT distribution over time. The analytic solution for the lower boundary choice probability (i.e. a NoGo response) is given by [28, 30, 31]:

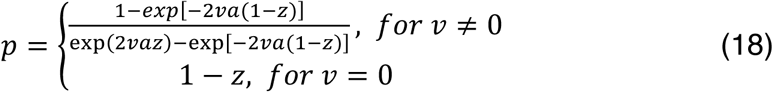

#### Model validation

The model variants were validated through prior predictive checks, parameter recovery analyses, and model identifiability assessments. Specifically, 20 synthetic datasets, each comprising data from 25 agents (500 agents total per model), were simulated using parameters sampled from the priors and hyperpriors from each model variant. The response time distributions and learning curves across all datasets were pooled and visualized to confirm that the models generated plausible behavioral data without a priori biases across experimental condition.

The synthetic behavioral datasets were subsequently re-fitted to their respective generating model variants for Bayesian parameter inference. To evaluate parameter recovery, Pearson correlation coefficients were calculated between the posterior means of the estimated parameters and the true data-generating values. Finally, to assess model identifiability, synthetic datasets generated by M1 and M2 were fitted to both model variants, and the Deviance Information Criteria (DIC) was computed for each model fit [32]. Model identifiability was evaluated by comparing the DIC values of the true generating model and the alternative model across datasets. Because M3 was an exploratory model developed post hoc based on the model comparison results of M1 and M2, it was omitted from the model identifiability analysis.

#### Model comparison

Model comparison between M1 and M2 was conducted both qualitatively, via posterior predictive checks, and quantitatively, using the DIC and Leave-One-Subject-Out Expected Log Predictive Density (LOSO-ELPD) [32, 33].

For posterior predictive checks, 8000 synthetic datasets were generated, each corresponding to a posterior sample and conditioned on the unique choice sequence assigned to each of the 25 participants. For each participant, simulated choices and response times were averaged across posterior draws to generate posterior predictive estimates of accuracy and response-time distributions across conditions, which were compared with the observed behavioral data. In addition, the 8000 synthetic datasets were used to construct 95% posterior predictive intervals for the pooled learning curves and response-time distributions across conditions, facilitating visual assessment of the models’ alignment with the observed data.

To quantitatively and comprehensively evaluate model performance, we used two complimentary metrics—the DIC and LOSO-ELPD—to assess in-sample fit and out-of-sample predictive accuracy, respectively. LOSO-ELPD was computed by repeating the model fitting procedure 25 times, leaving out the data of one participant per iteration. The resulting posterior draws were used to compute the expected log predictive density for each left-out participant i (ELPD_i_):

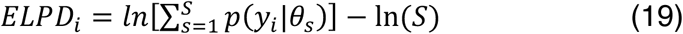

Where *y_i_* denotes the data of participant *i*, *θ_s_* denotes the posterior parameter draw obtained from fitting the model to the remaining 24 participants, and *S* denotes the total number of MCMC draws (*S*=8000). Each model viarant yielded 25 ELPD_i_ values, which were compared across models.

### Statistical Analyses

For statistical analyses involving behavioral measures (i.e. accuracy, response time), we adopted a conservative non-parametric approach due to the modest sample size (N = 25). Specifically, the Wilcoxon signed-rank test was used to compare behavioral measures across conditions, and Spearman’s rank correlation coefficient was computed to assess the associations between model parameters and behavioral measures. The significance level was set at α = 0.05, and Bonferroni correction was applied within each dataset according to the number of comparisons performed. For the parameter recovery assessment, Pearson’s correlation coefficient was used because the number of simulated agents was sufficiently large (N = 500). For Bayesian parameter estimates, we summarize posterior distributions using posterior means and 95% equal-tailed intervals (ETIs). Parameters with 95% ETIs excluding zero were interpreted as providing credible evidence for a non-zero effect. All statistical tests were conducted using the Python package scipy (version 1.17.1).

## Results

### Behavioral Analysis

We began by examining the behavioral measures: choice and response time. Fig. 3 shows the learning curves for the eight conditions averaged across 25 participants. By the end of the 15th trial, the proportion of Go responses exceeded 0.5 in all Go conditions and fell below 0.5 in all NoGo conditions, indicating appropriate learning at the group level.

**Figure 3:**
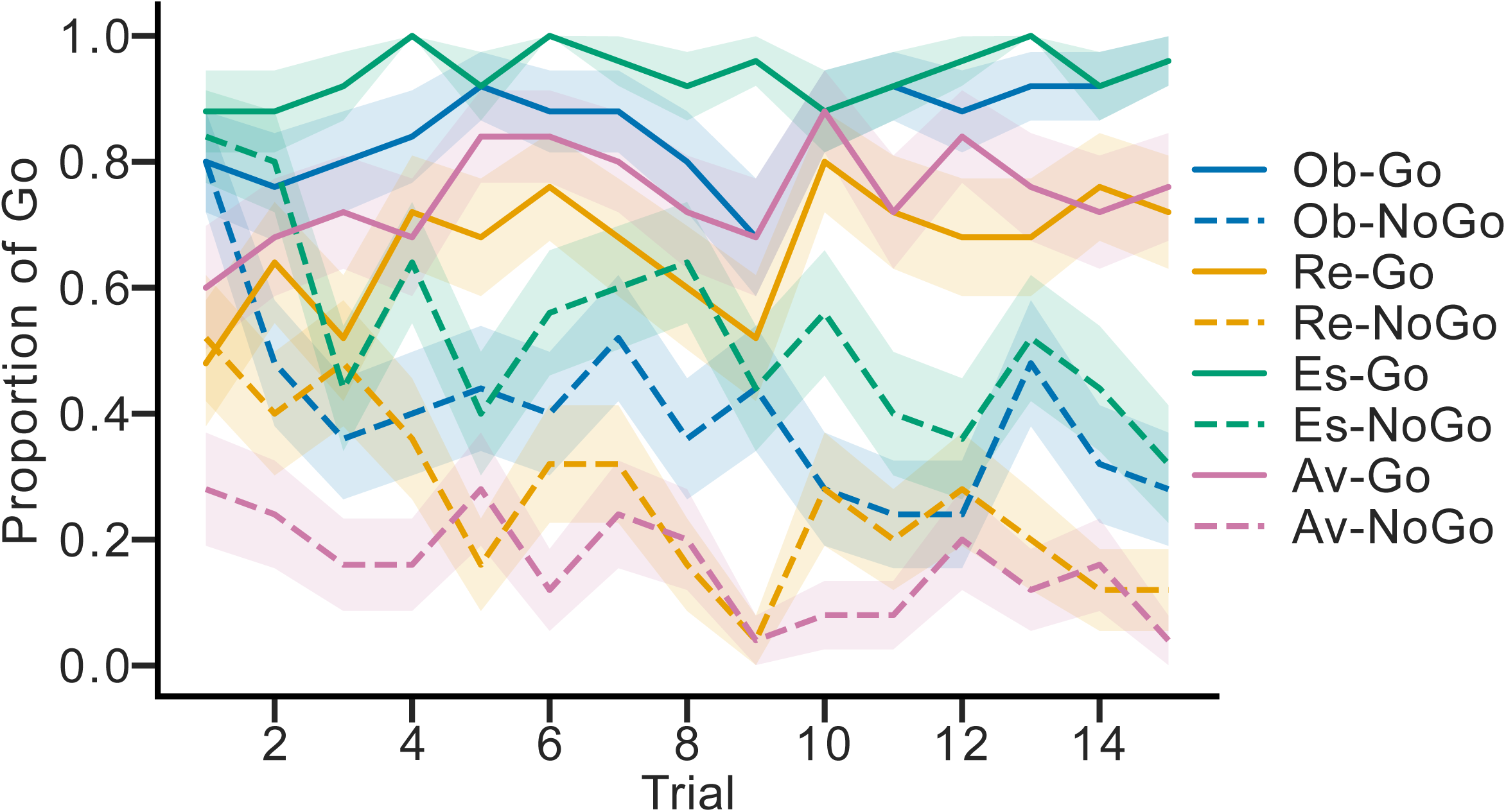
Condition-specific learning curves averaged across 25 participants. Curves depict the evolution of Go proportions across 15 trials. Conditions sharing the same cue type are indicated by color, with solid lines denoting Go conditions and dashed lines denoting NoGo conditions. Shaded areas represent ±1 standard error of the mean (SEM).

Several learning curves exhibited starting points that deviated from 0.5, suggesting the presence of group-level behavioral biases. Notably, conditions sharing the same cue (e.g., Obtain-Go and Obtain-NoGo) tended to begin at similar starting points, indicating that these biases were cue-specific. One notable exception was the Avoid cue, for which the starting points of the Go and NoGo learning curves diverged substantially more than those of the other cue types. This phenomenon is discussed further in the Discussion section. The mean initial proportion of Go responses for the four cue types was 0.80 for Obtain, 0.50 for Retain, 0.86 for Escape, and 0.44 for Avoid.

Fig. 4a shows the accuracy distributions for each condition. The left panel presents the raw block-wise accuracies of all 25 participants across the four task blocks, whereas the right panel further decomposes each block into its two constituent trial conditions (e.g., gain-congruent into Obtain-Go and Retain-NoGo, and gain-incongruent into Obtain-NoGo and Retain-Go), allowing condition-specific accuracy distributions to be examined. Here, accuracy was computed based on block-level correctness rather than trial-level correctness. For example, in the gain-congruent block, any Go response on an Obtain trial was counted as correct regardless of the feedback received on that particular trial.

**Figure 4:**
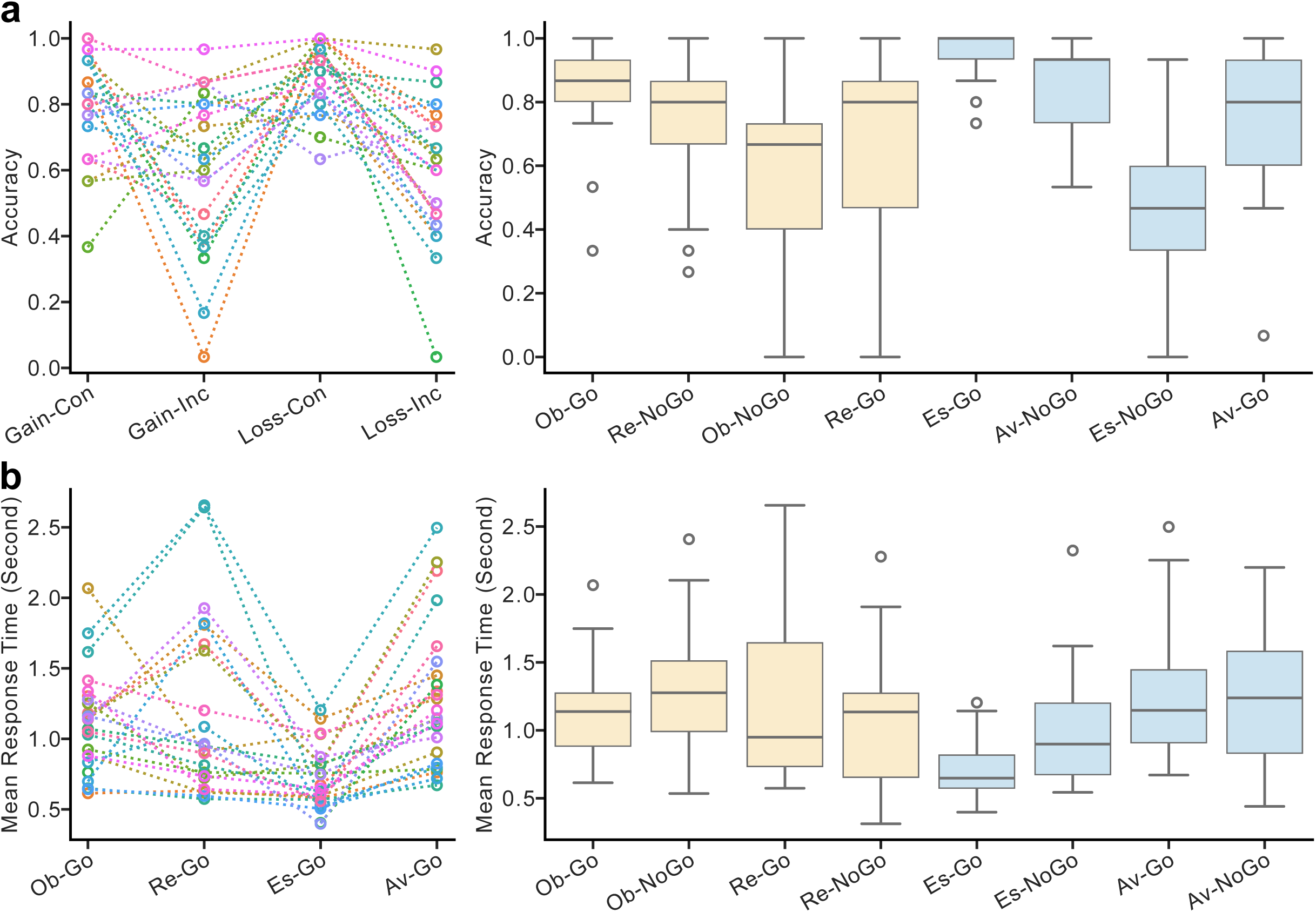
Distribution of condition-specific accuracy and mean response times across all participants. (a) Left panel: raw block-wise accuracy across all participants. Con: Congruent; Inc: Incongruent. Right panel: accuracy distributions across the eight conditions. (b) Left panel: mean response times for the four Go conditions. Right panel: mean response times across all eight conditions. Note that response times in NoGo conditions reflect incorrect Go responses and were excluded from primary analyses; they are included solely for completeness.

The left panel of Fig. 4a reveals a general trend of higher accuracy in action-congruent than action-incongruent blocks in both appetitive and aversive contexts (mean accuracies: gain-congruent 0.80; gain-incongruent 0.63; loss-congruent 0.89; loss-incongruent 0.61). When each block is further decomposed into its constituent trial conditions (Fig. 4a, right panel), the accuracy patterns across the four transtion-action combinations appear similar between the two valence contexts. However, the magnitude of the Go–NoGo mean difference is smaller in the appetitive context than in the aversive context (Obtain: 0.26; Retain: −0.07; Escape: 0.47; Avoid: −0.09). The directions of the Obtain, Escape, and Avoid differences were consistent with the findings of Guitart-Masip et al. and Millner et al. [12, 15]: the Obtain and Escape conditions exhibited a Go bias (i.e. Go – NoGo > 0), whereas the Avoid condition exhibited a NoGo bias (i.e. Go – NoGo < 0). Wilcoxon signed-rank tests were conducted across all four conditions, with Bonferroni correction applied (α = 0.0125). Significant Go–NoGo differences were observed only in the Obtain and Escape conditions (Obtain: p < 0.001; Retain: p = 0.64; Escape: p < 0.001; Avoid: p = 0.24).

Fig. 4b presents the mean response time distributions across experimental conditions. Because Go responses on NoGo trials represent incorrect responses that are more likely to be influenced by extraneous factors (e.g., attentional lapses, anticipatory responses, or exploratory behavior), the primary analyses focused on Go conditions (Fig. 4b, left panel), while the distributions across all conditions are shown in the right panel for completeness. On average, the response times are shorter in “change” conditions than in “stay” conditions within the same valence context (appetitive: Obtain-Go = 1.11 s, Retain-Go: 1.18 s; aversive: Escape-Go = 0.71 s, Avoid-Go = 1.29 s). This difference was larger and more consistent in the aversive context than in the appetitive context. Wilcoxon signed-rank tests with Bonferroni correction (α = 0.025) confirmed that the response-time difference between change and stay conditions was statistically significant in the aversive context (p < 0.001), but not in the appetitive context (p = 0.96). This behavioral pattern in the aversive context is consistent with the findings by Millner et al. [15], who similarly observed significantly faster RTs for Escape-Go than for Avoid-Go trials.

In summary, this novel behavioral paradigm successfully reproduced the key findings of Guitart-Masip et al. and Millner et al. [12, 15], including a Go bias in the Obtain and Escape conditions, a NoGo bias in the Avoid condition, and faster responses in the Escape than the Avoid condition. Moreover, the inclusion of the Retain condition enabled the evaluation of novel mechanisms involving action–transition congruency, complementing the existing valence-based framework.

### Rationale of Computational Models

The simplest explanation for the observed group-level bias patterns across the four trial structures is a transition-related bias modulated by context valence, with weaker effects in the appetitive context, as illustrated in Fig. 5b. However, closer inspection of the left panels of Fig. 4 suggests an alternative possibility: transition-related and salience-related biases offsetting each other in the appetitive context (Fig. 5c). This is based on the observation that the differences in accuracy and response time in the appetitive context do not merely decrease in magnitude; for some participants, they reverse direction (Fig. 4, left panels). This pattern is particularly evident in the mean-response-time distributions: Escape response times are consistently shorter than Avoid response times, whereas Retain response times appear to cluster into two subgroups—one longer than Obtain and the other shorter than Obtain. This appetitive-specific reversal, which reduces the apparent group-level effect, could be explained by individual differences in the relative strengths of transition-related and salience-related biases. As illustrated in Fig. 1 and 5c, the two biases operate in the same direction in aversive trial types but in opposite directions in appetitive trial types because the transition–cue pairings differ between the two contexts. Specifically, the non-neutral cue is paired with the “change” trial in the aversive context (i.e., Escape), whereas it is paired with the “stay” trial in the appetitive context (i.e., Retain). Consequently, when the transition-related bias is stronger than the salience-related bias (Fig. 5c, upper panel), the appetitive bias pattern resembles a weaker version of the aversive pattern because the salience effect partially offsets the transition effect. Conversely, when the salience-related bias exceeds the transition-related bias (Fig. 5c, lower panel), the appetitive bias pattern reverses because the salience effect outweighs the transition effect.

**Figure 5:**
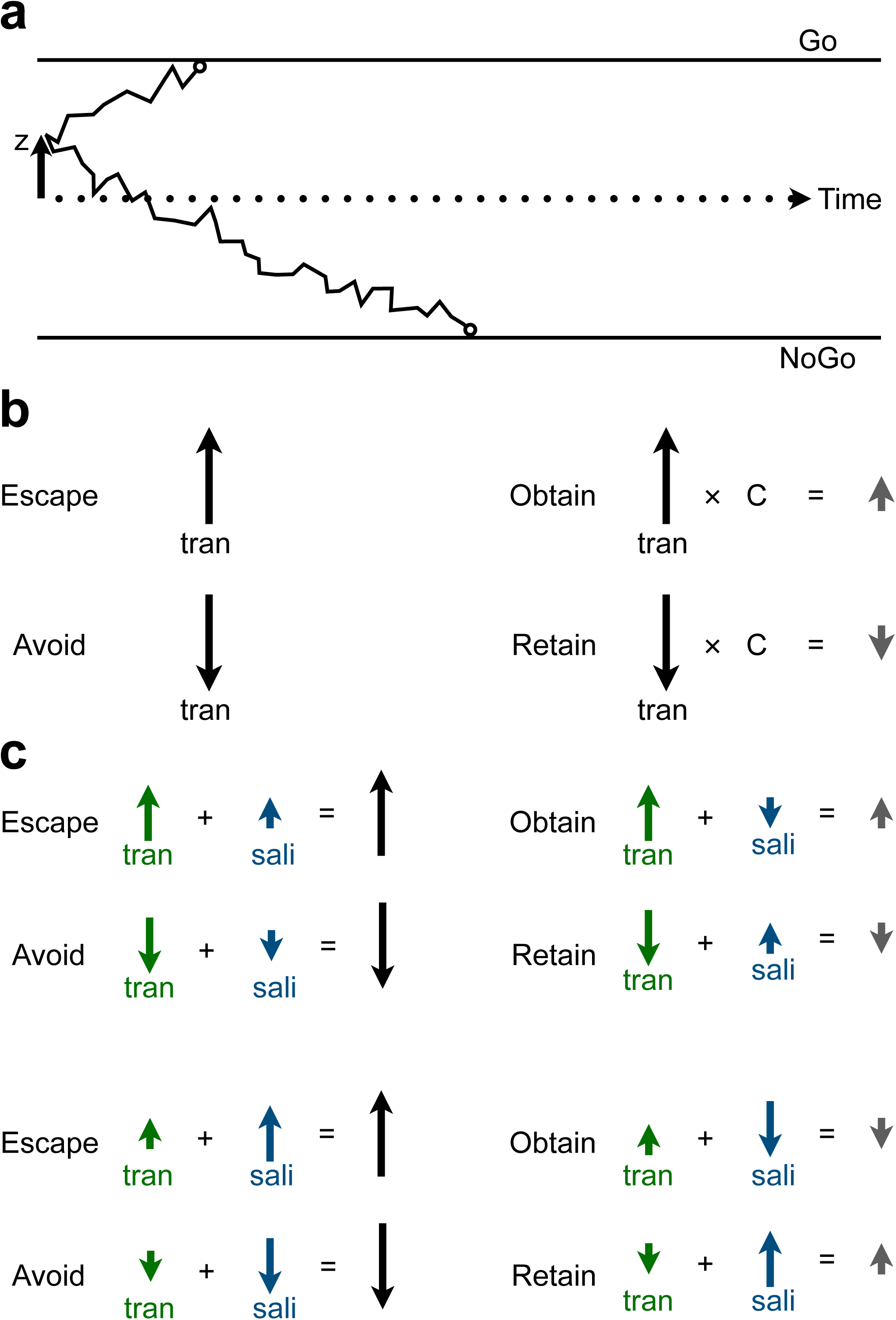
Conceptual diagram of the DDM framework and competing mechanistic interpretations. (a) Basic schematic of a DDM. A decision process is conceptualized as a drift process where a “decision variable” originates at starting point *z* (arrowhead, left) and drifts at rate *V* according to a stochastic Wiener diffusion process until reaching one of two decision boundaries (i.e., Go vs. NoGo). The sign and magnitude of *V* is determined by the difference of the Go and NoGo Q-values. The *z* parameter influences both choice probability and decision latency independently of *V* by setting a baseline choice bias. (b) The context–valence modulation model, in which attenuated bias in appetitive contexts is attributed to valence-dependent scaling. C denotes the scaling factor for transition-related bias in the appetitive context. (c) The transition–salience offsetting model, in which attenuated bias in appetitive contexts arises from mutually opposing transition-related and salience-related biases. Green arrows denote transition-related bias, blue arrows denote salience related bias.

If the transition–salience offsetting model illustrated in Fig. 5c is correct, the reversals observed in Fig. 4 should reflect a structured pattern rather than random noise, with their direction determined by the relative magnitudes of the two biases. This prediction can be tested using computational modeling. Importantly, this approach addresses a key limitation highlighted by Millner et al. [15], whose paradigm could not dissociate bias driven by cue non-neutrality from biases arising from other sources (e.g. two-factor theory).

### Computational Modelling

#### Model Validation

Two hierarchical Bayesian RL-DDM models, M1 and M2, were constructed to represent the transition–salience offsetting and context-valence modulation hypotheses, respectively. To evaluate parameter interpretability and model identifiability, 20 synthetic behavioral datasets, each comprising 25 agents, were simulated from each model and subsequently cross-fitted to both models.

Synthetic datasets generated by M1 and M2 were pooled separately to examine prior predictive response-time distributions and condition-specific learning curves (Supplementary Fig. 1a and 1b). The simulated response-time distributions for both models, with incorrect responses represented as negative response times, fell within a plausible range (−3 to 3 s), with correct responses occurring more frequently than incorrect responses. Mean learning curves across all conditions initiated at approximately 0.5, reflecting the null hypothesis that no condition-specific biases were present. By the end of the 15th trial, all learning curves approached approximately 0.8 or 0.2, consistent with appropriate learning of the 80%/20% task contingency.

Parameter interpretability was evaluated by assessing the models’ capacity to recover the true data-generating parameters through self-fitting (Supplementary Fig. 2a, 2b, 3a, 3b). All parameters were recovered to an acceptable level (Pearson correlation coefficient r > 0.75), with the exception of *cxt_dz* in M2 (r = 0.68). This reduced recoverability reflects an intrinsic limitation arising from the interaction between a multiplicative factor and a prior centered at zero. Because *cxt_dz* scales *tran_dz*, it can be reliably recovered only when *tran_dz* is sufficiently large. This consideration was taken into account when interpreting the model-fitting results presented in the following section (Supplementary Fig. 4).

To assess model identifiability, each of the 20 synthetic datasets generated by M1 and M2 was fitted to both its true generating model and the alternative model, with DIC values calculated for self-fits and cross-fits. Supplementary Fig. 5 presents the resulting confusion matrix, displaying the proportion of datasets identified by each model based on lower DIC values. M1 correctly recovered 90% of M1-generated datasets, whereas M2 correctly recovered 60% of M2-generated datasets. That M1 was favored in 40% of M2-generated datasets aligns with M2 being functionally nested within M1; specifically, datasets generated under M2 can satisfy M1’s condition wherein transition-related bias outweighs salience-related bias (Fig. 5b and the top panel of Fig. 5c).

#### Model Fitting

Having confirmed satisfactory model performance and interpretability, we proceeded to empirical model fitting. Fig. 6 presents the fitting results of M1 and M2 for the experimental data. As shown in Fig. 6a, both the M1 posteriors of the group-level transition-related bias and salience-related bias (*tran_dz_μ* and *sali_dz_μ*, respectively) substantially deviated from the zero-centered informative priors. The posterior mean of *tran_dz*_*μ* was 0.34 (95% ETI = [0.23, 0.46]), whereas the posterior mean of *sali_dz*_*μ* was 0.25 (95% ETI = [0.16, 0.34]). Neither ETI overlaps with zero. These values strongly indicate a group-level Go-bias associated with both the “change” transition and cue non-neutrality.

**Figure 6:**
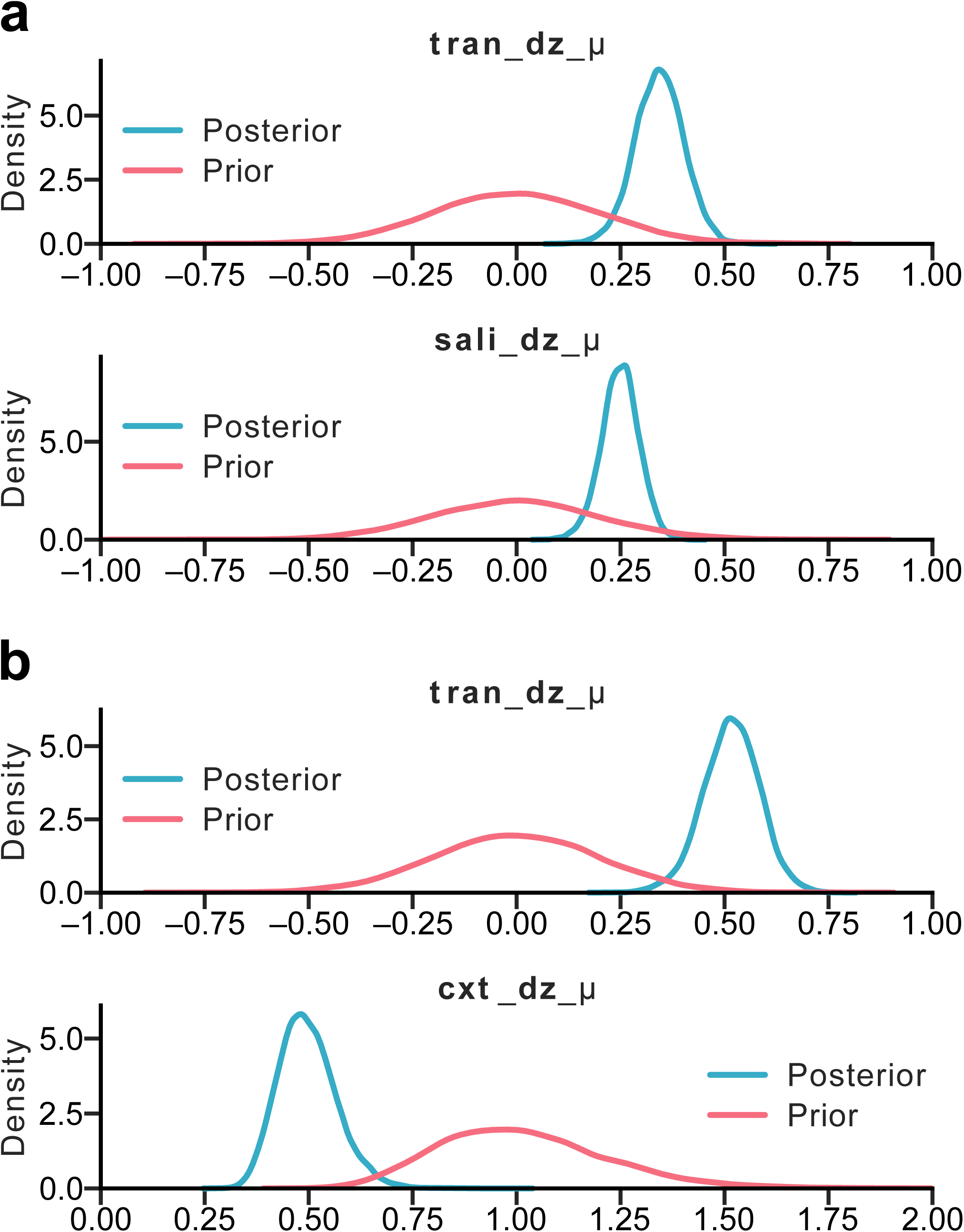
Model fitting results for M1 and M2. (a) Prior and posterior distributions for the group-level transition-related bias parameter (*tran_dz_μ*) and salience-related bias parameter (*sali_dz_μ*) in model M1. (b) Prior and posterior distributions for the group-level transition-related bias parameter (*tran_dz_μ*) and context-valence modulation parameter (*cxt_dz_μ*) in model M2.

Fig. 6b displays the group-level priors and posteriors for M2 (i.e. *tran_dz*_*μ* and *cxt_dz*_*μ*). Similar to M1, M2 identified a pronounced Go-bias associated with the “change” transition, with a posterior mean of *tran_dz*_*μ* of 0.52 (95% ETI = [0.38, 0.65]). The posterior of *cxt_dz*_*μ* shifted leftward relative to its log-normal prior centered at one, yielding a posterior mean of 0.50 (95% ETI = [0.38, 0.65]). Values below 1 indicate an attenuated transition-related bias in appetitive contexts, aligning with empirical behavioral observations. All *z*-modulation parameters across M1 and M2 demonstrated satisfactory MCMC chain convergence (R^ < 1.01; ESS > 400).

As discussed in the parameter recovery analysis, context–valence modulation in M2 can be reliably estimated only when transition-related bias is sufficiently large. Given the robust magnitude of transition-related bias in M2, the estimate of context–valence modulation can be considered highly reliable. For reference, the distribution of individual *tran_dz* estimates across all 25 participants is provided in Supplementary Fig. 4.

#### Model Comparison

The model-fitting results indicate that both M1 and M2—representing two competing hypotheses—captured meaningful condition-specific modulations underlying the observed bias patterns. To qualitatively compare M1 and M2, 8000 datasets were simulated from MCMC posterior draws to construct 95% posterior predictive intervals (PPIs) for all group-level learning curves and response-time distributions (Supplementary Fig. 6a and 6b). The 95% PPIs from both models overlapped well with experimental observations. Simulated datasets were also averaged at the participant level to derive accuracy and response-time distributions across conditions, matching the format of the left panels of Fig. 4. As illustrated in Fig. 7, both M1 and M2 captured the general pattern of reduced accuracy in incongruent blocks and shorter mean response times on “change” trials; however, accuracy and mean-response-time reversals were exhibited exclusively in M1 simulations.

**Figure 7:**
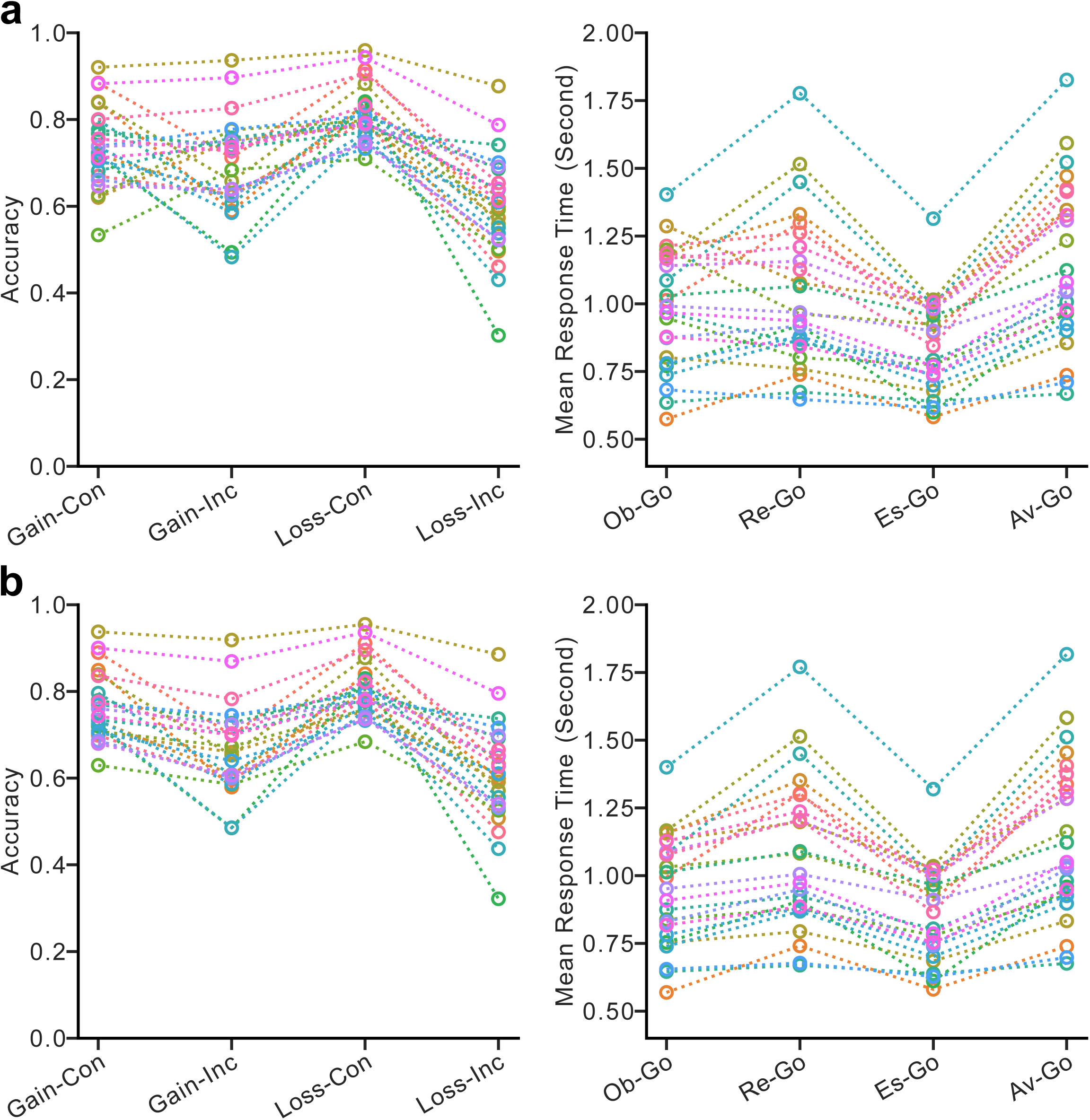
Posterior predictive distributions of block-wise accuracy and condition-specific mean response times. Data generated from posterior samples across all participants for (a) model M1 and (b) model M2.

For quantitative comparison, DIC values were calculated for both models. M1 yielded a lower DIC than M2 (M1: 4386.07; M2: 4439.58), reflecting superior in-sample fit. To evaluate out-of-sample predictive performance, LOSO-ELPD was computed for each participant and compared across models. The distribution of participant-level M1– M2 LOSO-ELPD differences is presented in Supplementary Fig. 7. This differece averaged 1.05 ± 2.52 (mean ± standard deviation), indicating a numerically higher predictive performance for M1, although this difference was not statistically significant according to the Wilcoxon signed-rank test (p = 0.29).

#### Model Prediction of Behavior

The model comparison results indicate that, although both M1 and M2 capture group-level behavioral patterns satisfactorily, M1 may provide a more nuanced fit at the individual level. To further evaluate the empirical implications of this difference, we examined whether the parameter estimates from M1 and M2 predicted individual-level behavioral observations in ways consistent with the patterns illustrated in Fig. 5. Specifically, under the transition–salience offsetting hypothesis (M1; Fig. 5c), behavioral biases, as captured by accuracy differences between congruent and incongruent blocks and reflected in mean response times, should correlate with *tran_dz* – *sali_dz* in appetitive contexts and *tran_dz* + *sali_dz* in aversive contexts. Conversely, the context–valence modulation hypothesis (M2; Fig. 5b) predicts that such biases should correlate with *tran_dz* ✕ *cxt_dz* in appetitive contexts and with *tran_dz* in aversive contexts. Furthermore, the ratio of these composite parameters across valence contexts in the two models should predict individual-level differences in relative bias magnitude.

Fig. 8 illustrates the correlations between congruent–incongruent accuracy differences and composite model parameters across valence contexts, alongside cross-context ratio relationships, for both model variants. Consistent with theoretical predictions, both models showed significant positive correlations in Spearman’s rank correlation analyses with Bonferroni correction (α = 0.025). Crucially, in M1, the linear relationship within the appetitive context (Fig. 8a, left and right panels) spans both positive and negative values, thereby accounting for empirical accuracy reversals. This continuum indicates that, rather than reflecting random noise, negative reversals are governed by the same offsetting mechanism driving positive bias values—specifically, the dominance of salience-related bias over transition-related bias. In contrast, corresponding M2 prediction (Fig. 8b, left and right panels) shows a marked clustering of data points at near-zero values, highlighting M2’s inability to account for these reversal dynamics.

**Figure 8:**
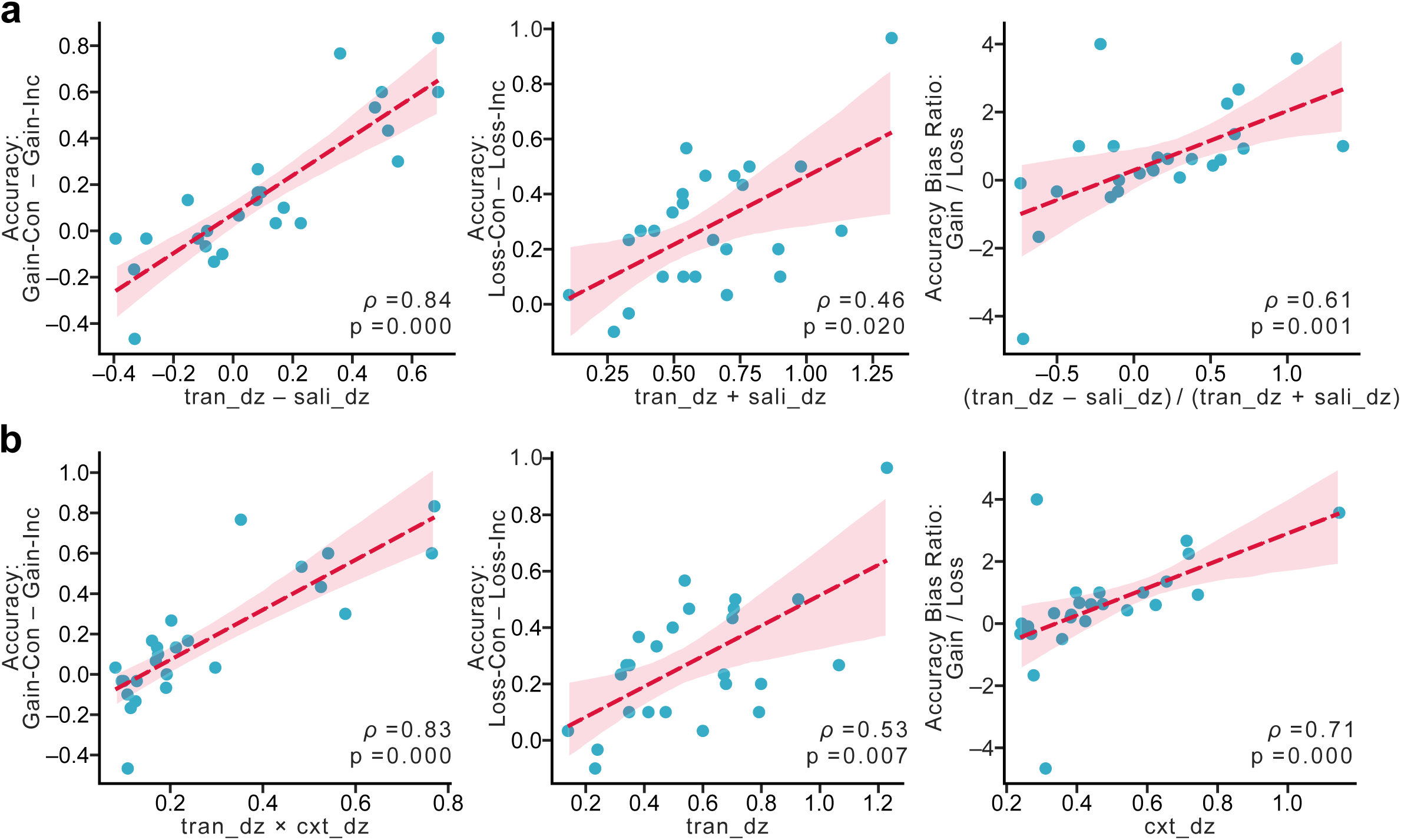
Participant-specific parameters predict accuracy bias. (a) Correlations between composite parameters formed by *tran_dz* and *sali_dz* and accuracy bias patterns across valence contexts, as predicted by model M1. (b) Correlations between composite parameters formed by *tran_dz* and *cxt_dz* and accuracy bias patterns across contexts, as predicted by model M2. Spearman’s rank correlation coefficients and p-values are reported in the lower-right corner of each panel.

We also examined the relationships between composite parameters and mean response-time differences across valence contexts (Fig. 9). Although response-time data carry lower statistical sensitivity than accuracy metrics owing to the exclusion of all block-level incorrect trials and NoGo responses in Go conditions, both models captured weak, non-significant negative trends between composite parameters and response-time differences, aligning with model predictions. Notably, M2 again exhibited a moderate clustering of data points near zero in the appetitive context (Fig. 9b, left), a pattern absent in M1 (Fig. 9a. left). This pattern further underscores M2’s inability to account for response-time reversals that are effectively captured by M1.

**Figure 9:**
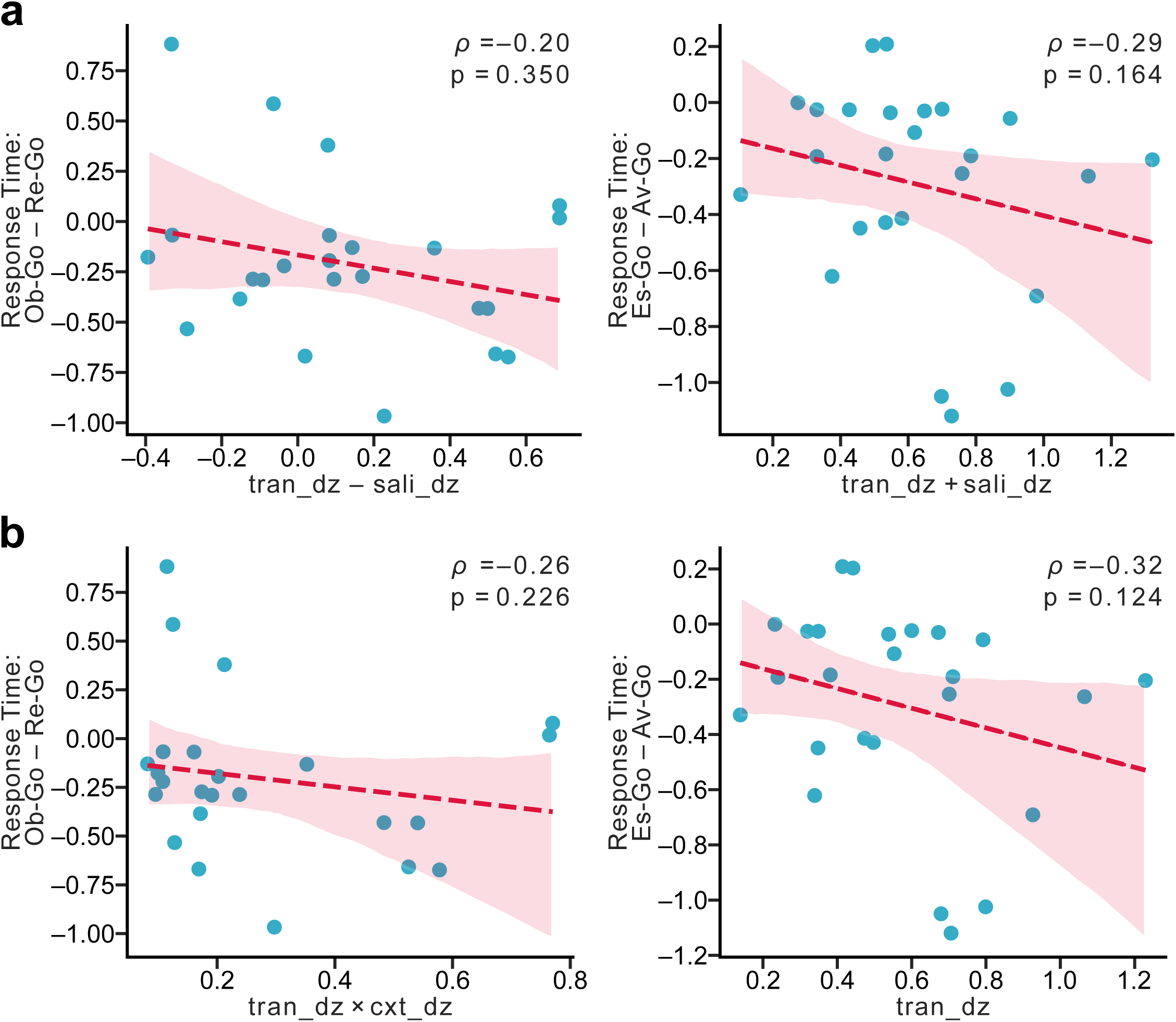
Participant-specific parameters predict response-time bias. (a) Correlations between composite parameters formed by *tran_dz* and *sali_dz* and response-time bias patterns across valence contexts, as predicted by model M1. (b) Correlations between composite parameters formed by *tran_dz* and *cxt_dz* and response-time bias patterns across valence contexts, as predicted by model M2. Spearman’s rank correlation coefficients and p-values are reported in the upper-right corner of each panel.

#### Exploratory Model

The findings described above indicate that M1, which lacks an explicit valence component, explains the experimental data better than M2, which relies on valence modulation. This observation may seem surprising given that Pavlovian bias, which orthogonal Go/NoGo paradigms were originally designed to isolate [12, 15], is defined as motor biases driven by emotionally-valence contexts. To investigate whether contextual valence plays any role in shaping the observed behavioral patterns, we constructed M3, extending the winning model M1 by incorporating a context-valence-dependent bias modulator (*cxt_dz*). Unlike in M2, *cxt_dz* in M3 isolates context-valence effects independently of *tran_dz*. To align with the theory proposed by Guitart-Masip et al. that appetitive stimuli promote Go responses while aversive stimuli facilitate NoGo responses [12], *cxt_dz* is assigned a positive sign in the appetitive context and a negative sign in the aversive context (equation (13)-(16) in Methods). M3 was rigorously validated through the same evaluation pipeline (i.e., prior predictive checks and parameter recovery analyses; Supplementary Fig. 1c, 2c, and 3c) prior to empirical model fitting.

Fig. 10a presents the model fitting results. Similar to M1, M3 yielded robust positive estimates for *tran_dz*_*μ* and *sali_dz*_*μ*, with posterior means of 0.34 (95% ETI = [0.21, 0.46]) and 0.21 (95% ETI = [0.10, 0.31]), respectively. Notably, *cxt_dz*_*μ* was estimated to be negative rather than positive, with a posterior mean of –0.14 (95% ETI = [-0.23, –0.034]). This finding indicates that, contrary to the interpretation of Guitart-Masip et al. [12], the aversive context promotes Go responses rather than NoGo responses. Model fit was evaluated through qualitative posterior predictive checks following the procedures used for M1 and M2 (Supplementary Fig. 6c). Condition-specific accuracy and mean-response-time distributions simulated from 8000 posterior draws are displayed in Fig. 10b. Like M1, M3 accounts for condition-specific motor biases while capturing accuracy and response-time reversals. M3 achieved a DIC of 4299.30, lower than that of M1, demonstrating superior in-sample fit. For out-of-sample predictive performance, the M3– M1 LOSO-ELPD difference averaged 0.83 ± 3.22 (mean ± standard deviation). The Wilcoxon signed-rank test indicated no significant difference in LOSO-ELPD between M1 and M3 (p = 0.89).

**Figure 10:**
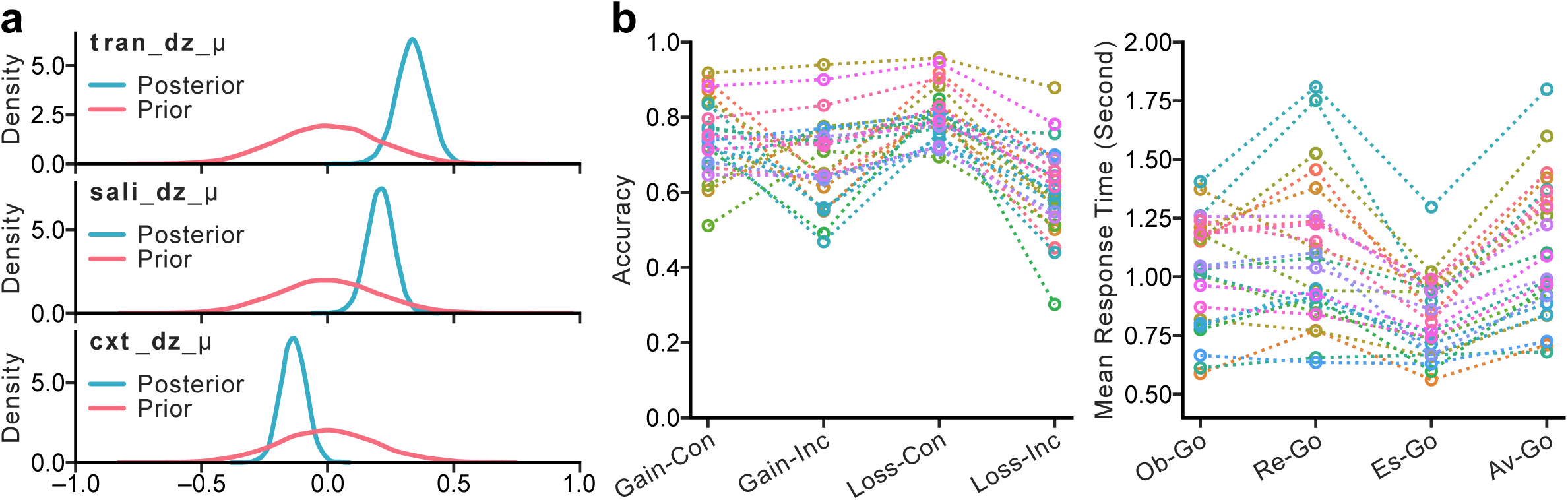
Model fitting results and posterior predictive distributions for model M3. (a) Prior and posterior distributions for the group-level transition-related bias (*tran_dz_μ*), salience-related bias (*sali_dz_μ*), and context–valence modulation (*cxt_dz_μ*) parameters in model M3. (b) Posterior predictive distributions of block-wise accuracy and condition-specific mean response times generated from posterior samples.

## Discussion

The present study used a novel orthogonal valenced Go/NoGo paradigm to decompose Pavlovian bias—the motor bias traditionally thought to be driven by emotionally valenced contexts—into putative non-emotional components, thereby providing crucial new insights into classic findings from orthogonal Go/NoGo paradigms and their translational relevance for psychiatric research [11, 12, 15, 18]. In addition to successfully replicating the key findings of previous studies—a Go bias for obtaining rewards and escaping threats, a NoGo bias for avoiding punishment, and shorter response times for escaping than for avoiding punishment—the novel Retain condition revealed a comparatively mild NoGo bias with more heterogeneous behavioral patterns. Building on these findings, we propose that the bias isolated by orthogonal paradigms primarily reflects an association between “action” and “change”, and between “inaction” and “staying unchanged”, rather than an intrinsic effect of emotional valence. To account for the attenuated bias observed in the appetitive context, we evaluated two context-dependent modulation mechanisms: transition–salience offsetting and valence-dependent scaling. Hierarchical RL-DDM analyses favored the transition–salience offsetting model, which explains the observed behavioral patterns without requiring an emotional-valence component. An exploratory extension of this winning model further suggested that, beyond aimed transition and cue salience, context valence exerts a secondary influence by invigorating action across both change and stay trials in the aversive context rather than serving as the primary driver of the observed bias patterns.

These findings address several issues from previous studies that were either unrecognized or left unresolved. The first concerns the mismatch between the aversive NoGo bias reported by Guitart-Masip et al. and the fight-or-flight account of threat responding, which predicts hyperarousal and action invigoration under stress [12–15]. As illustrated in Fig. 1, the reward and punishment conditions (i.e. Obtain and Avoid) used to establish the appetitive–aversive contrast in Guitart-Masip et al. was confounded by a change–stay contrast. Based on our behavioral findings and RL-DDM modeling results, this transition-related contrast exerts an effect opposite to that of the appetitive–aversive contrast and may dominate the observed bias pattern under the original experimental design. By fully orthogonalizing these two contrasts, our paradigm dissociates their respective contributions, providing a cleaner assessment of the independent effects of transition and context valence on motor biases. The same interpretation extends to the bias patterns in the aversive context reported by Millner et al [15, 18]. In their paradigm, participants aimed to change the state during Escape trials but to maintain the current state during Avoid trials. Our modeling results suggest that this difference in aimed transition can directly account for the observed Go–NoGo bias without invoking the valence reversal required by the two-factor theory.

An additional consideration in the work of Millner et al. is the presence of non-neutral cues in the escape trials, which further complicates the interpretation of the observed bias patterns [15]. Millner et al. acknowledged that cue non-neutrality—equated to “aversiveness” in their discussion—could itself contribute to the Go bias, but their task was unable to disentangle this possibility from other putative mechanisms [15]. Our paradigm isolates this factor from a broader perspective by evaluating the effect of cue non-neutrality (i.e., salience) across oppositely valenced contexts, rather than treating it as a property unique to aversive stimuli. This distinction is important because salience may invigorate action through increased arousal regardless of emotional valence [20–22]. Using the same RL-DDM framework adopted by Millner et al., our analyses successfully isolated an action-invigorating effect of salience across oppositely valenced contexts, supporting the hypothesis that cue salience contributes to the Go bias independently of aimed transition and context valence. These modelling results align with the observations of Millner et al., but suggest a more nuanced interpretation: the Go bias attributed to “aversiveness” may instead reflect a more general effect of cue salience.

Aside from the theoretical clarifications, the findings of the present study raise a broader empirical question regarding clinical construct validity: what psychological process does the “Pavlovian bias” isolated by orthogonal Go/NoGo paradigms actually reflect, and how does it mechanistically relate to clinical symptoms? Because Pavlovian bias has long been conceptualized as a motor bias driven by emotional contexts, findings from these paradigms are often interpreted as reflecting aberrant emotional processes. For example, in Millner et al., the active escape bias was interpreted as a heightened desire to escape from pain and linked to suicidal behavior [18]. Our findings, however, suggest that the motor biases observed in valenced orthogonal Go/NoGo paradigms are largely accounted for by non-emotional components that operate across both appetitive and aversive contexts. Although conducted in a normative sample, our study—with classic findings replicated and previously conflated factors dissociated—invites a reconsideration of the mechanistic interpretation linking Pavlovian bias to clinical symptoms, as well as the underlying neural dynamics that give rise to these behavioral effects [34].

Another consideration for the ecological validity of orthogonal Go/NoGo paradigms, briefly discussed by Millner et al. [15], concerns the choice of sensory modality for valenced stimuli. While Guitart-Masip et al. followed the convention of monetary incentivization [12], Millner et al. sought to better emulate the “psychological pain” associated with psychiatric disorders by adopting a primary sensory modality—unpleasant auditory noise [15, 18]. Although diversifying task designs improve generalizability, several considerations regarding data reliability, construct interpretability, and clinical applicability ultimately led us to adopt a valence scheme closer to that of Guitart-Masip et al. First, primary sensory modalities (e.g., auditory, olfactory, or tactile stimuli) are susceptible to sensory adaptation and habituation [35–38], which can substantially alter task performance even within a single experimental session without careful calibration. Second, while a Go response to a primary aversive stimulus can be interpreted within a goal-directed framework (i.e., to “escape”), responses to primary pleasant stimuli are inherently more difficult to interpret. In the context of auditory stimulation, for example, both a lullaby and a dance song can be classified as pleasant stimuli, yet they would be expected to influence motor behavior in opposite ways. Third, many psychiatric disorders are characterized by atypical sensory processing, potentially confounding task performance independently of the cognitive mechanisms of interest. For example, autistic spectrum disorder is frequently associated with auditory and tactile hypersensitivity, as well as altered sensory habituation, which may complicate data interpretation and raise ethical concerns about the use of unpleasant primary sensory stimulation [39–41]. To balance these considerations, we adopted secondary appetitive and aversive stimuli that lacked direct real-world financial connotations (e.g., avoiding dollar signs, coins, and banknotes) and provided a fixed monetary compensation that was independent of task performance. This design preserves the advantages of secondary reinforcement while reducing both associations with personal financial circumstances and sensory-specific confounds, making it more broadly applicable across clinical populations. Consistent with this rationale, our task successfully replicated the key behavioral findings reported by both Guitart-Masip et al. and Millner et al. [12, 15], suggesting that it captures the same core psychological processes while providing greater flexibility for translational research.

Some limitations—and opportunities for improvement—warrant further discussion. As shown in Fig. 3, the starting points of the two Avoid learning curves diverged substantially more than those of the other three cue types. Both curves began closer to their respective plateau values, yet their average starting point remained slightly below 0.5. This pattern suggests that additional learning processes beyond those captured by the current RL-DDM may have influenced early task performance. One plausible explanation is the combined effect of cross-trial learning and valence-dependent learning rates. Because each block contained only two trial types (e.g., Obtain-Go and Retain-NoGo in the gain-congruent block), some participants may have implicitly learned that when one trial type was associated with a Go response, the other was more likely to be associated with a NoGo response, despite this relationship not being explicitly stated in the task instructions. This cross-trial learning may have been particularly pronounced in the Avoid condition because learning rates are often higher in aversive contexts, as reported in previous studies [42, 43], and because the average starting point of Avoid trials was closer to 0.5, providing greater room for learning-related shifts during the early trials. Given that this was a validation study with a modest sample size, we adopted the simplest modeling framework needed to test our primary hypotheses. Consequently, we did not incorporate additional mechanisms, such as cross-trial learning or valence-dependent learning rates, which may have led to suboptimal model fit, particularly in the aversive context. Future studies employing similar paradigms should consider explicitly integrating these mechanisms, as our findings suggest they may play a meaningful role in driving decision dynamics.

In conclusion, the present study dissociated several factors that can independently contribute to motor biases but have previously been conflated in Pavlovian bias research. Our findings suggest that the motor bias patterns isolated by orthogonal Go/NoGo paradigms are primarily accounted for by two non-emotional components: aimed transition and cue salience. Given the widespread application of Pavlovian bias paradigms in research on affective disorders and suicidality, these findings call for a careful reconsideration of how Pavlovian bias is conceptualized in clinical research and how findings from these paradigms are mechanistically interpreted.

## Data Availability

Per institutional ethical requirements, de-identified behavioral data are available from the corresponding author upon reasonable request only for participants who provided explicit consent for their data to be retained and shared. Data from participants who did not provide such consent will not be shared. Aggregated data and simulated data supporting the findings of this study are available from the corresponding author upon reasonable request.

## Code Availability

The code used for computational modelling, data processing, and statistical analyses is available from the corresponding author upon reasonable request.

