## Supplementary Information for "Active Escape or Active Change? Decomposing Pavlovian Bias into Non-emotional Components"

#### Part I: RL-DDM priors

The priors and hyperpriors of the hierarchical Bayesian RL-DDM parameters common to all three model variants are specified below. The relative starting point (*z*) was further modulated by the condition-specific z-modulators (*tran_dz*, *sali_dz*, *cxt_dz*) as described in the Methods section of the main text. The non-decision time parameter (*t*) was estimated at the subject level to accommodate individual differences in minimal response time (*min_RT_per_subject*), which sets an upper bound on allowable non-decision time. Trials with response times shorter than 0.2 s were considered invalid and excluded prior to model estimation.

<u>Learning rate (*alpha*):</u>

$$alpha\_\mu\_logit \sim Normal(0,0.6)$$

$$alpha\_\sigma \sim HalfNormal(0.4)$$

$$alpha\_offset \sim Normal(0, 1)$$

$$alpha = sigmoid(alpha\_\mu\_logit + alpha\_offset \times alpha\_\sigma)$$

<u>Boundary separation (*a*):</u>

$$a\_\mu\_log \sim Normal(0.3,0.29)$$

$$a\_\sigma \sim HalfNormal(0.25)$$

$$a\_offset \sim Normal(0,1)$$

$$a = exp(a\_\mu\_log + a\_offset \times a\_\sigma)$$

Relative starting point (*z*):

$$z\_\mu\_logit \sim Normal(0,0.4)$$

$$z\_\sigma \sim HalfNormal(0.3)$$

$$z\_offset \sim Normal(0,1)$$

$$z = sigmoid(z\_\mu\_logit + z\_offset \times z\_\sigma)$$

Transition-related z-modulator (M1, M2 and M3):

$$tran\_dz\_\mu \sim Normal(0,0.2)$$

$$tran\_dz\_\sigma \sim HalfNormal(0.2)$$

$$tran\_dz\_offset \sim Normal(0,1)$$

$$tran\_dz = tran\_dz\_\mu + tran\_dz\_offset \times tran\_dz\_\sigma$$

Salience-related z-modulator (M1 and M3):

$$sali\_dz\_\mu \sim Normal(0,0.2)$$

$$sali\_dz\_\sigma \sim HalfNormal(0.2)$$

$$sali\_dz\_offset \sim Normal(0,1)$$

$$sali\_dz = sali\_dz\_\mu + sali\_dz\_offset \times sali\_dz\_\sigma$$

Context-valence z-modulator of M2:

$$cxt\_dz\_\mu\_log \sim Normal(0{,}0.2)$$

$$cxt\_dz\_\sigma \sim HalfNormal(0.2)$$

$$cxt\_dz\_offset \sim Normal(0{,}1)$$

$$cxt\_dz = exp(cxt\_dz\_\mu\_log + cxt\_dz\_offset \times cxt\_dz\_\sigma)$$

Context-valence z-modulator of M3:

$$cxt\_dz\_\mu \sim Normal(0{,}0.2)$$

$$cxt\_dz\_\sigma \sim HalfNormal(0.2)$$

$$cxt\_dz\_offset \sim Normal(0{,}1)$$

$$cxt\_dz = cxt\_dz\_\mu + cxt\_dz\_offset \times cxt\_dz\_\sigma$$

Scaling factor for Q-value difference (*scaler*):

$$scaler\_\mu\_log \sim Normal(0.89{,}0.25)$$

$$scaler\_\sigma \sim HalfNormal(0.3)$$

$$scaler\_offset \sim Normal(0{,}1)$$

$$scaler = exp(scaler\_\mu\_log + scaler\_offset \times scaler\_\sigma)$$

Non-decision time (*t*):

$$t\_frac\_\mu \sim Beta(2{,}2)$$

$$t\_frac\_\kappa \sim Beta(4{,}2)$$

$$t\_frac \sim Beta[t\_frac\_\mu \times t\_frac\_\kappa, (1 - t\_frac\_\mu) \times t\_frac\_\kappa]$$

$$t = 0.2 + t\_frac \times (\min\_RT\_per\_subject - 0.01 - 0.2)$$

### Part II: Supplementary Figures

**Supplementary Figure 1: Prior predictive checks for the three model variants.**

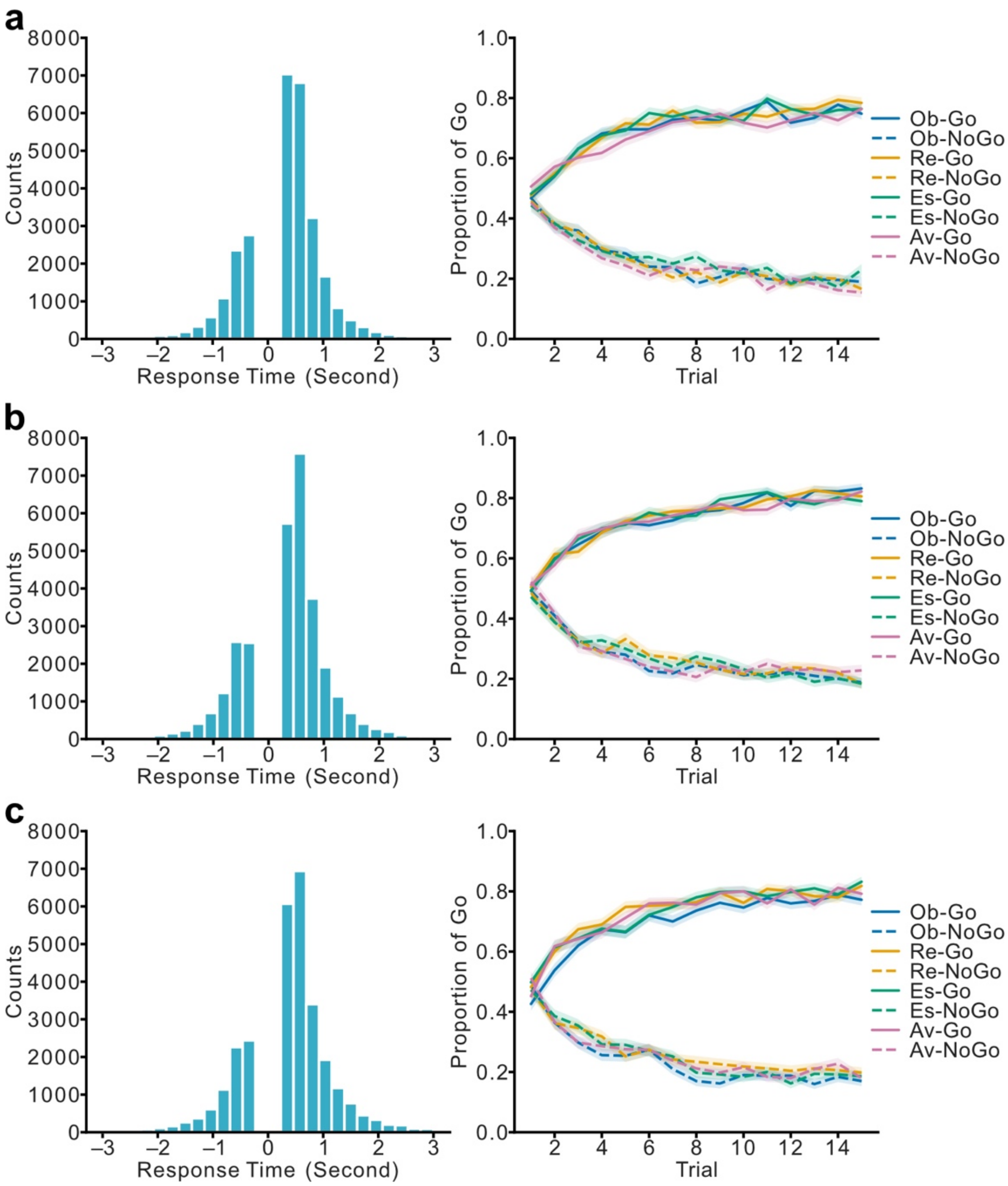


(a) M1, (b) M2, and (c) M3. Left panels display response time distributions pooled across 20 simulated datasets (25 agents per dataset) for each model, with block-level correct and incorrect responses represented as positive and negative values, respectively. Right panels show condition-specific average learning curves across the eight conditions over

15 trials derived from the simulated datasets; shaded areas indicate the standard error of the mean (SEM).

**Supplementary Figure 2: Group-level parameter recovery assessment of the three model variants.**

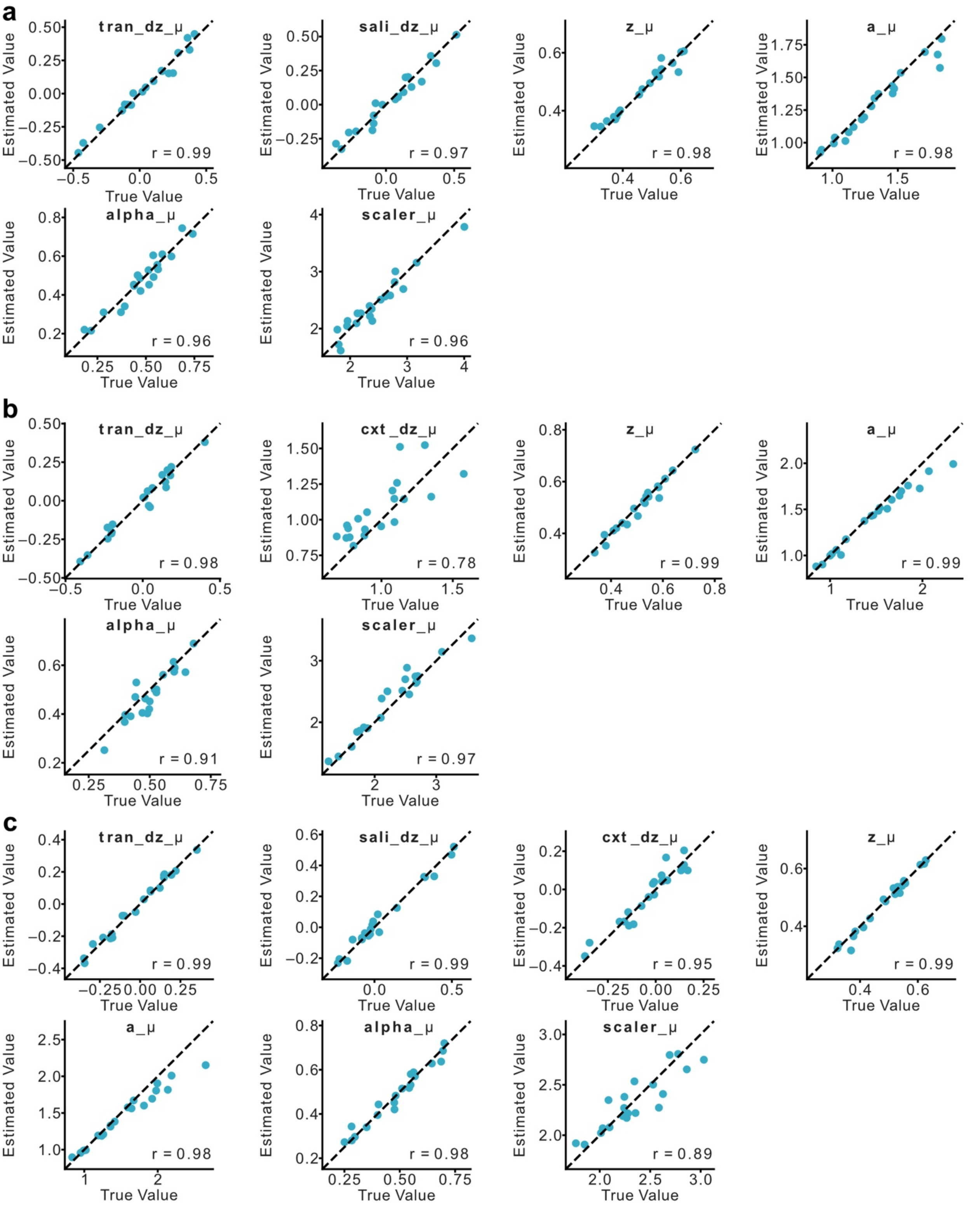

(a) M1, (b) M2, and (c) M3. Scatter plots display the relationship between true data-generating parameters and estimated parameters across 20 simulated datasets, with the corresponding Pearson correlation coefficients shown in the lower right of each panel. For ease of interpretation, parameters estimated in unconstrained spaces were transformed back to their respective bounded spaces as specified in the RL-DDM framework (e.g. *z_μ_logit* transformed to *z_μ* through the sigmoid transform).

**Supplementary Figure 3: Individual-level parameter recovery assessment for the three model variants.**

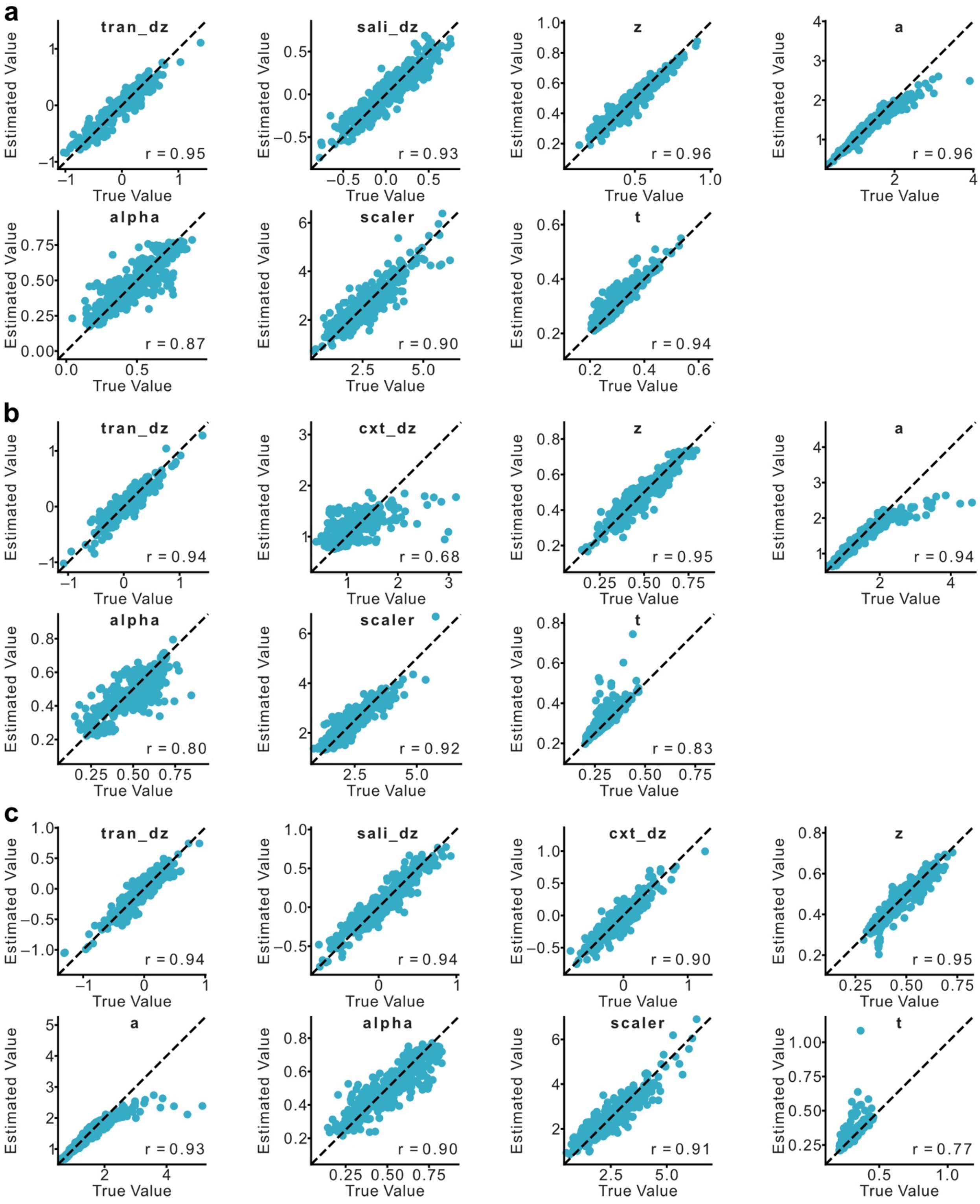

(a) M1, (b) M2, and (c) M3. Scatter plots display the relationship between true data-generating parameters and estimated parameters across all 500 individual agents comprising the 20 simulated datasets. Pearson correlation coefficients are reported in the lower right of each panel.

**Supplementary Figure 4: Parameter interpretability constraint of *cxt_dz* in model M2.**

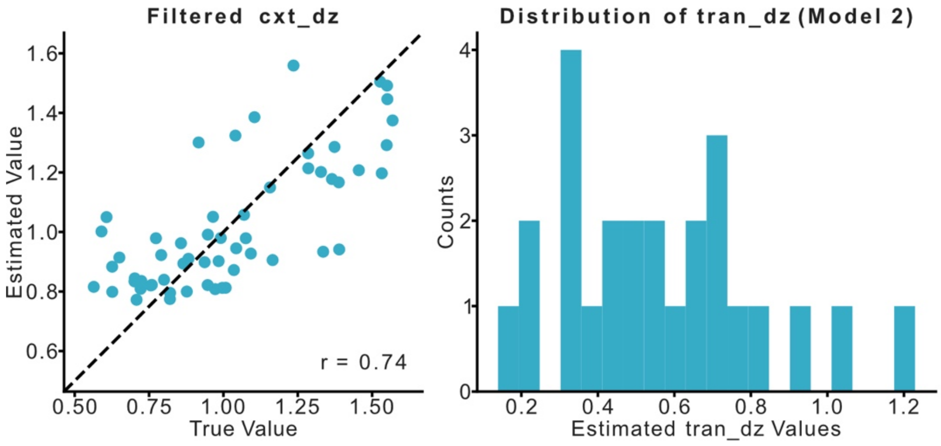


Left panel: The recovery of *cxt_dz* improves (Pearson's r increases from 0.68 to 0.74) when data points with *tran_dz* < 0.35 are excluded, illustrating how the magnitude of *tran_dz* impacts the recoverability of *cxt_dz* due to the mathematical formulation of M2. Right panel: Empirical distribution of *tran_dz* estimated from the experimental dataset.

**Supplementary Figure 5: Model identifiability confusion matrix for M1 and M2.**

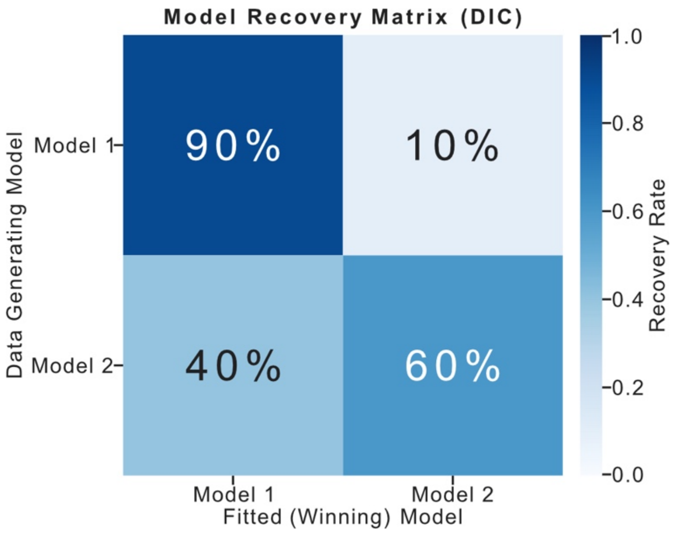


M1 correctly recovered 18 out of 20 self-generated datasets (90% recovery rate), whereas M2 recovered 12 out of 20 self-generated datasets (60% recovery rate). Successful recovery was defined as the true data-generating model achieving a lower Deviance Information Criterion (DIC) value than the alternative model.

**Supplementary Figure 6: Qualitative posterior predictive checks.**

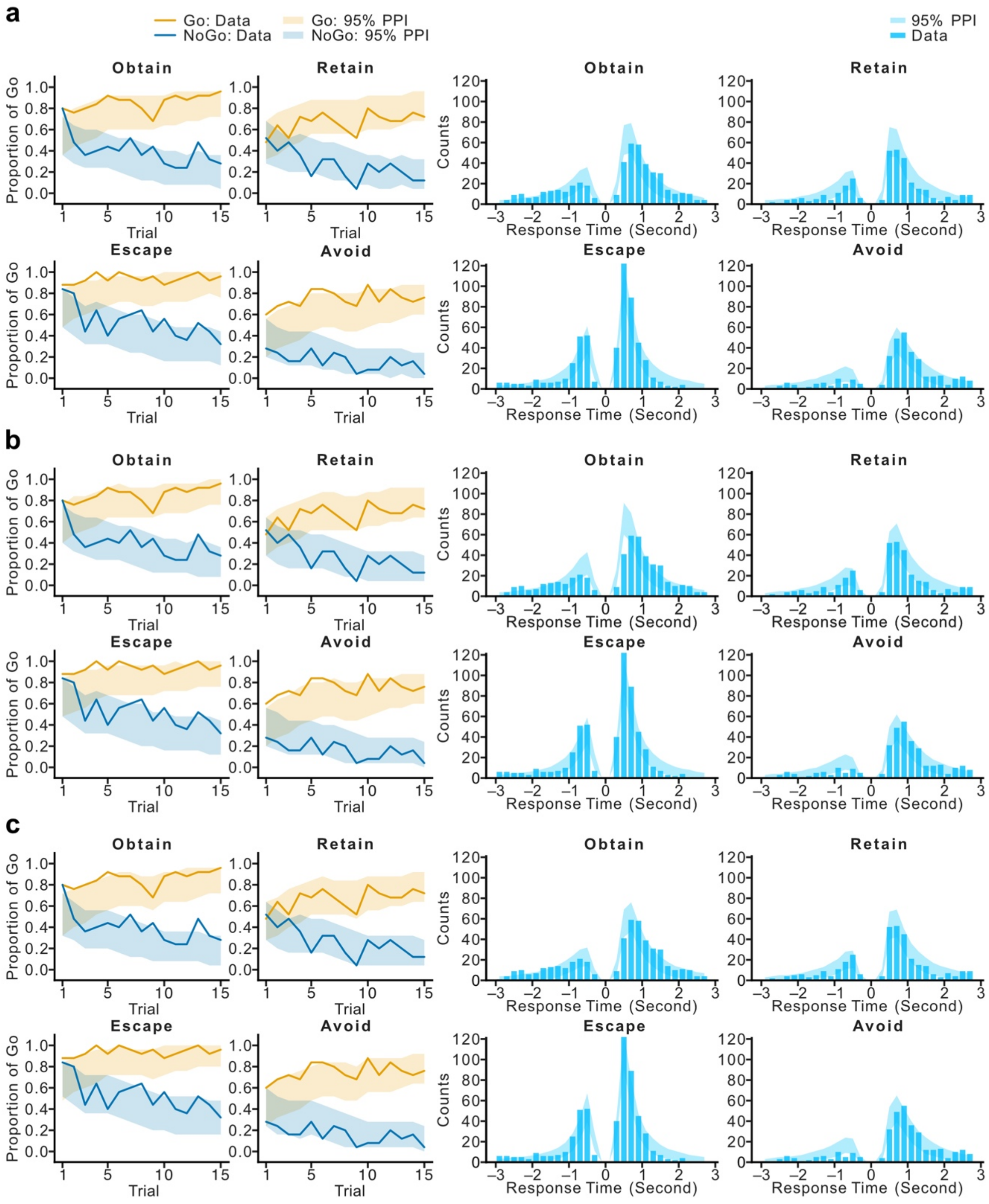

(a) M1, (b) M2, and (c) M3. To enhance visual clarity, empirical learning curves across the eight conditions were grouped by cue type and plotted across four subfigures. The 95% posterior predictive intervals derived from the modeling results were overlaid to evaluate alignment between model predictions and empirical data. Similarly, 95% posterior predictive intervals for condition-specific response time distributions were overlaid on empirical distributions for visual comparison.

**Supplementary Figure 7: The distribution of the differences of LOSO-ELPD between M1 and M2 across 25 participants.**

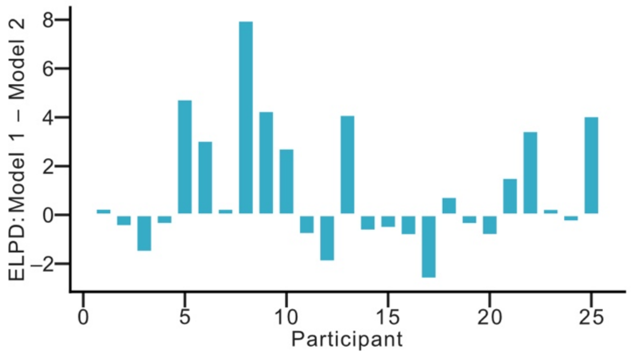
